# Social compensation changes predicted effects of individual loss in animal societies

**DOI:** 10.64898/2026.11.18.767549

**Authors:** Adelaide D. Abraham, Eloise F. Newman, Ben C. Sheldon, Sarah C. L. Knowles, Josh A. Firth

## Abstract

Animal populations globally are suffering significant declines; many species are social, and individuals are likely to adjust their social behaviour as they lose associates. Therefore, social networks may respond to population decline as individuals adjust their social connections in response to changing social environments. Our understanding of how individual responses to loss of their associates alters the structure of their social networks is rudimentary. To quantify how social responses to individual loss can alter network structure, we simulated individual loss and compensatory formation of new social ties across 268 empirical animal social networks. Compensation changed the structural effects of individual loss, though these effects often depended on which individuals were removed. Loss of central, peripheral or random individuals produced different network responses, while social compensation had similar effects regardless of compensation strategy. Together, these results suggest that compensatory social behaviour can reorganise social networks as populations decline, even if departing from the original structure. Accounting for these responses may improve predictions of how demographic change affects social structure, particularly when losses are socially patterned and without knowing the compensation strategy. Broadly, our findings highlight compensation as a potentially important process linking individual social behaviour to the reorganisation of animal societies.

## Main Text

Animal populations globally are declining under anthropogenic pressure (Loss, Will, and Marra 2015; Hill, DeVault and Belant 2019, Spooner et al. 2018). The loss of individuals from a population can have important consequences for the social structure of animal societies, as captured through social networks (Shizuka and Johnson 2020, Jarić et al. 2026). The likelihood of individual loss can depend on an individual’s social network position within the social network, such as socially peripheral individuals experiencing higher mortality (Ellis et al. 2017; Stanton and Mann 2012, Williams and Lusseau 2006). Individual animals are unlikely to be passive in response to such changes, for example, individuals may shift their social associations in response to losing associates (Firth et al. 2017; Downing et al. 2023). These individual behaviours determine network structure, yet how individual responses to loss shape network structure remains poorly understood, particularly across real-world animal systems.

When individuals are lost from a network, remaining individuals may respond by forming new associations or strengthening existing ones. Such reorganisation of social connections is often described as network rewiring (Han and Goetz 2019; Watts and Strogatz 1998). Here, we use ‘compensation’ to refer to rewiring that occurs after node loss, where remaining individuals adjust their social connections following the loss of associates. Evidence from animal systems suggests that individuals can respond socially after the loss of associates, but relatively little is known about how such responses alter network structure across species, removal contexts and compensation strategies (Shizuka and Johnson 2020; see Meier et al. 2025 for a discussion of cultural responses to node loss). If individuals respond to disturbance by recovering their preferred social network position (Gartland et al. 2022), these new associations may help stabilise population-level network structure while individual social environments change (Farine 2021). Alternatively, compensatory change could exaggerate the effects of the loss alone (Weber et al. 2013). Despite this, studies of population decline rarely account for the social responses of surviving individuals, and compensation remains poorly understood across animal social networks.

Because patterns of individual loss and social compensation likely differ among populations, species and contexts, a broad comparative approach is needed to understand when compensation may alter the network consequences of demographic change. Existing work has largely examined compensation in theoretical or simplified network settings with limited comparison across animal social systems (Artime et al. 2024). Here, we test how simulated individual loss and compensatory formation of new social ties alter network structure across 268 real-world animal social networks. We compare three forms of node removal with three simple compensation algorithms, alongside a no-compensation control, to ask whether compensation changes the structural consequences of individual loss and whether this varies with the pattern of loss or the compensation strategy (Figure 1).

**Figure 1:**
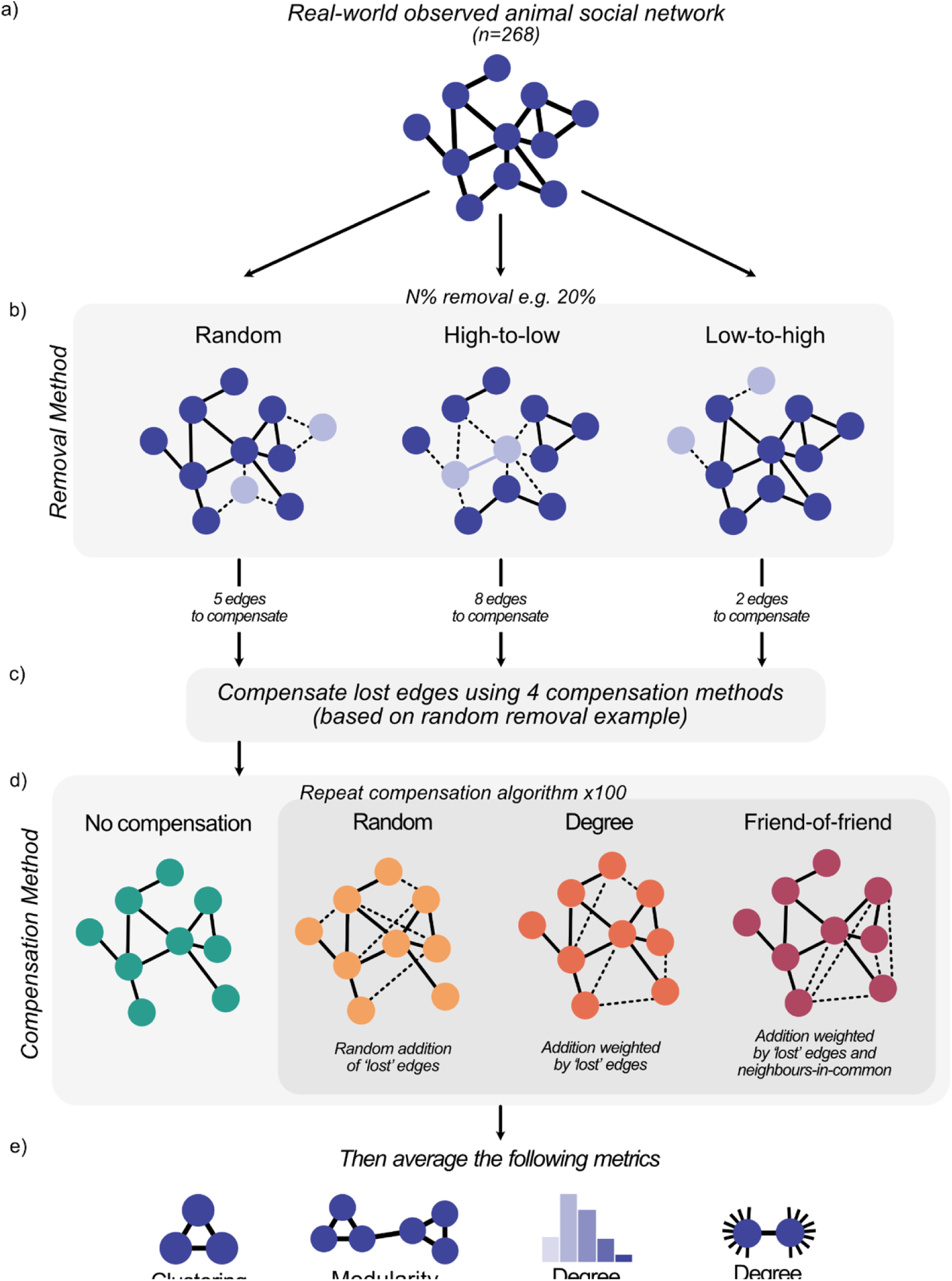
Conceptual figure of overall methodology of simulation. 268 animal social networks follow the following procedure: (a) different percentages of individuals (nodes) are removed from the network using three removal methods (b). Three compensation strategies (c) are then compared with a no-compensation control (d). For the compensation strategies, the number of edges added into the network is equal to the number of edges lost by the remaining individuals who were previously connected to individuals which were removed. For each compensation method this is repeated 100 times. The average of four network metrics (e) is calculated for each simulated network and averaged across compensation replicates. Clustering is the tendency of individuals to form local clusters. Modularity is the extent a network can be divided into distinct groups or communities. Degree heterogeneity is the variation in the number of connections of individuals in the network. Degree assortativity represents the correlation between the degrees of connected nodes. The whole pipeline is repeated separately for 10%, 20% and 30% node loss scenarios on each original network.

## Results

We simulated the effect of removal in 268 animal social networks, at three different removal percentages (10%, 20%, 30%), with three different removal methods (random, high-to-low centrality, and low-to-high centrality). We then simulated compensation of lost edges within the network, either randomly, by biasing new ties towards individuals that lost associates (i.e. prioritising maintenance of degree), or by prioritising forming connections with friends of friends.

### Removal method has predictable impacts on network structure

In the absence of compensation, the effects of node removal on network structure varied depending on removal method (Figure 2). Random removal did not significantly change clustering, degree heterogeneity or degree assortativity, but did significantly increase modularity only at 30% removal of nodes (Supplementary Information Figure SI.18). High-to-low removal significantly increased modularity and degree heterogeneity (Figure SI.18 and Figure SI.19), significantly decreased clustering (Figure SI.17), but did not change degree assortativity (Figure SI.20). In comparison, low-to-high removal decreased all metrics except clustering, which significantly increased (Figure SI.17).

**Figure 2:**
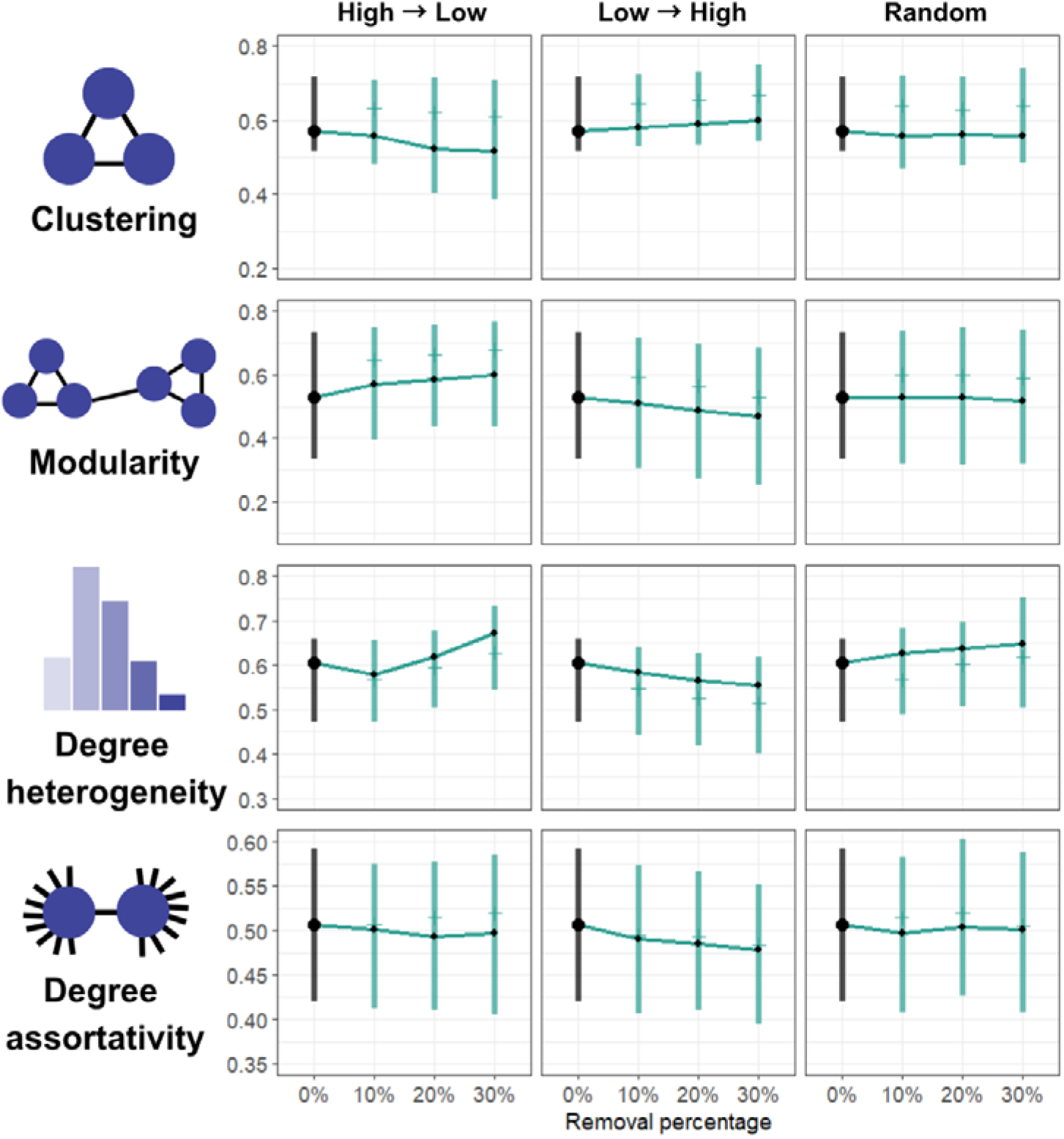
Comparison of network metrics for three removal methods; high-low centrality, low-high centrality and random removal. As all networks started at the same metric values, metrics at step 0 are coloured black. Black points represent mean values, while the line connecting points tracks means across each distinct removal percentage. Line ranges contain 50% of the data, between the 25% and 75% quantile. Small side bars represent median values.

### Presence of compensation, rather than type, shows different patterns

Compensation after node loss significantly altered network structure compared to no compensation for all four metrics (Figure 3). Notably, all compensation methods caused a decrease in network metric compared to node removal without compensation. Relative to their pre-removal values, the only metrics to not show a significant change with compensation were clustering under low-to-high removal and degree heterogeneity with high-to-low removal and friend-of-friend compensation (Figure 3; Figure SI.17 and Figure SI.19).

**Figure 3:**
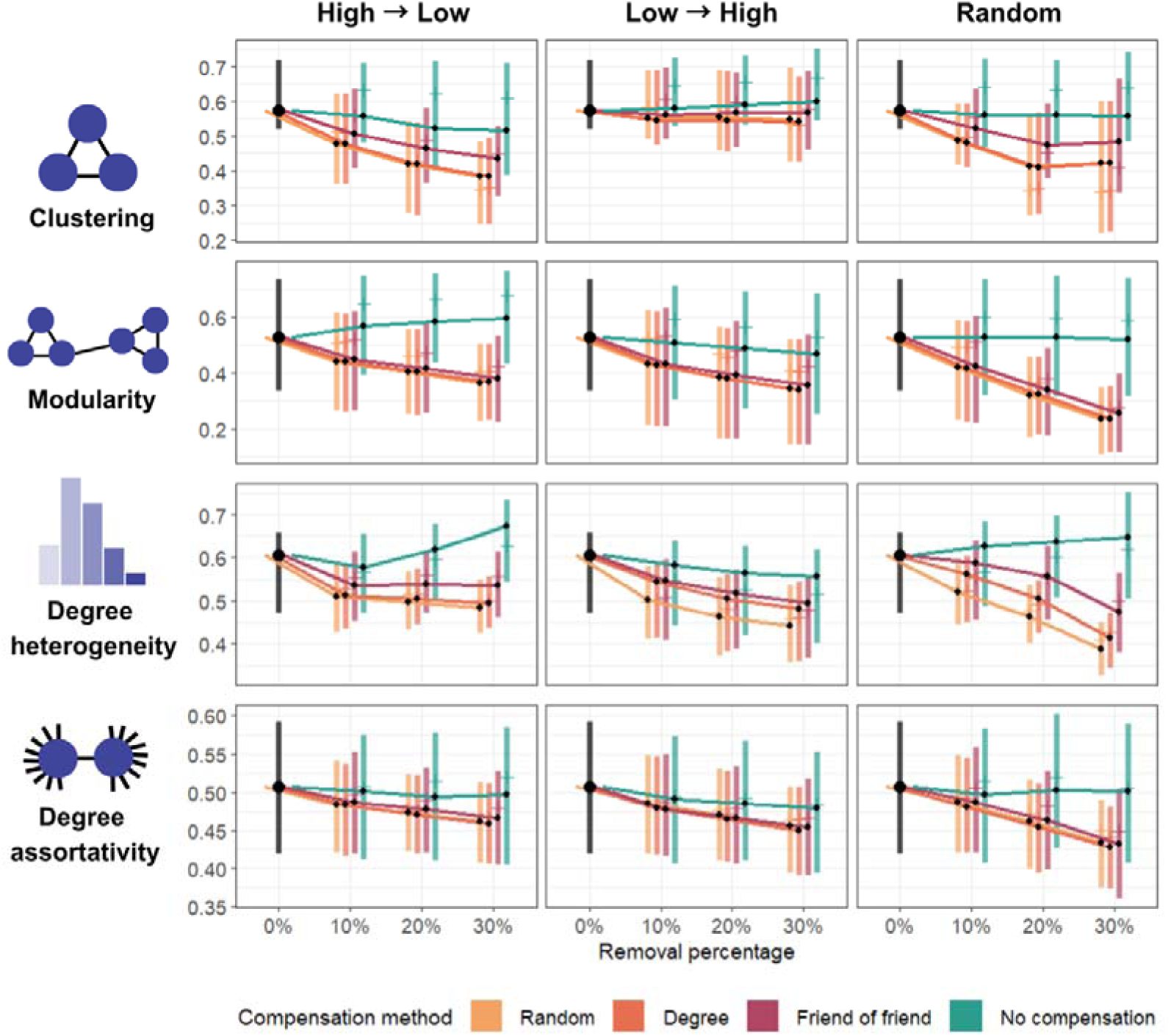
Comparison of network metrics for three removal methods; high-low centrality, low-high centrality and random removal. Compensation method is shown with no compensation (teal), random compensation (yellow), degree-biased compensation (orange), and bias to friends of friends (red). As all treatments share the same set of pre-removal networks, metrics at step 0 are coloured black. Black points represent mean values, while the line connecting points tracks averages across each distinct removal percentage. Line ranges contain 50% of the data, between the 25% and 75% quantile. Small side bars represent median values.

The tested compensation methods produced broadly similar patterns in the four network metrics, with the presence of compensation seeming more important than the specific type of compensation. Friend-of-friend clustering was significantly different from other compensation methods across all removal types (Figure SI.9). There were no significant differences in modularity between compensation methods, with the exception of friend-of-friend and degree compensation being significantly different at 30% low-to-high removal (Figure SI.10). At 30% removal, all compensation methods had significantly different degree heterogeneity with random removal, while friend-of-friend and degree did not differ under low-to-high removal, and random and degree did not differ under high-to-low (Figure SI.11). There were no significant differences in degree assortativity between compensation methods (Figure SI.12).

### Removal and compensation effects on network structure persist after accounting for phylogeny

These trends occurred in the context of large amounts of variation between networks, in both starting value and response. We modelled log multiplication factor (the scale of change in metric value between 0% and 30% removal) for each metric, accounting for the phylogenetic relationships between species as one of the sources of this variation. Removal method, compensation method, their interaction, and start value of the metric were included as fixed effects. Effect estimates (Figure 4) showed similar results as described above from the raw values (Figure 3), showing that removal method and the presence of compensation are both important for predicting network structural change, but that specific type of compensation method does not matter. The only effect not observed after account for phylogeny was the effect of low-to-high or high-to-low removal on clustering, suggesting that differences in clustering between removal methods can be explained by network phylogeny (95% credible interval *low to high:* −0.010 – 0.076*; high to low:* −0.064 – 0.035; note that this effect was present in a model without random effects (Figure SI.24)). Most estimated effects were retained when phylogenetic covariance was included in the model (Figure 4) and therefore are not explained by phylogenetic relationships between species included here. Additionally, these results show how the initial network structure (‘start value’) may predict the scale of metric change for clustering, degree assortativity, and degree heterogeneity (*clustering:* −0.205 – −0.439; *degree heterogeneity:* −0.370 – −0.269; *degree assortativity*: −1.280 – −1.157).

**Figure 4:**
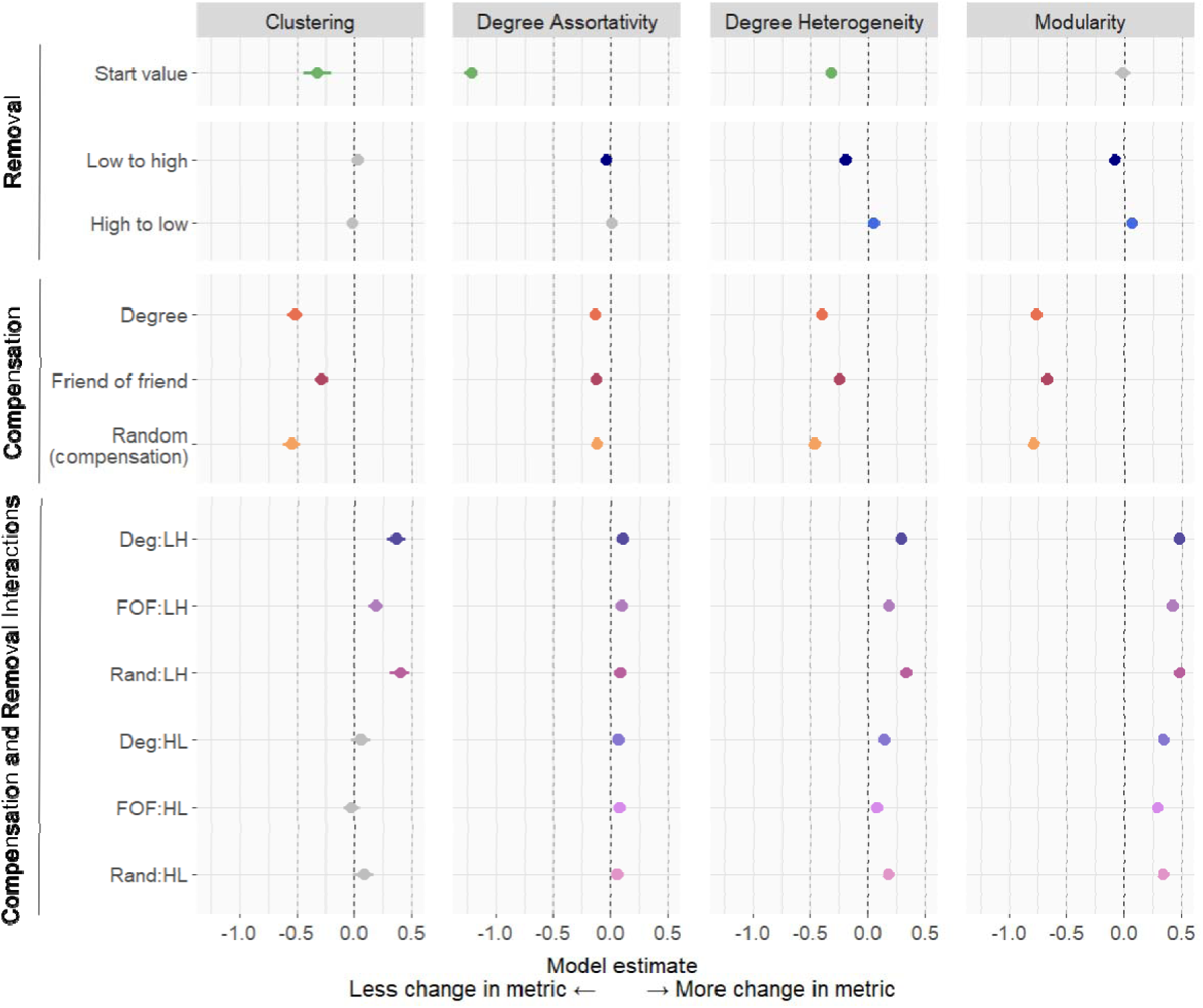
Estimated effect of removal method, compensation method, and start value on the multiplication factor of each metric, after accounting for phylogenetic pseudoreplication. Estimates are calculated using a Bayesian Student’s t model in brms (Bürkner 2021). Points are point estimates, with lines representing 95% credible intervals. Random removal and no compensation are used as reference levels. Acronyms are as follows: Deg (degree compensation); FOF (Friend of friend compensation); Rand (Random compensation); LH (Low-to-high removal); HL (High-to-low removal). Points coloured in grey have credible intervals that include zero.

## Discussion

Our findings suggest that the network consequences of individual loss depend on both the pattern of loss and the potential for social compensation. The loss of individuals altered network structure, with different effects depending on whether individuals were removed randomly, from high-to-low centrality or low-to-high centrality. Allowing remaining individuals to compensate for loss by forming new social ties changed all four network metrics, showing that social behaviour in response to loss can modify the structural impacts of individual loss, even if the networks are not restored to their original state. This pattern is still observed after accounting for phylogenetic relationships between network species, meaning that our findings are not only explained by taxonomic similarities between networks. These results suggest that predicting how social networks respond to demographic change may require attention to who is lost and whether remaining individuals adjust their associations.

In animal populations, mortality, emigration and managed removals are unlikely to be random with respect to social position, and many factors (e.g. age, sex, dominance, health, behaviour) can covary with network position. Therefore, demographic turnover - among other processes - may remove some kinds of social individuals more often than others (Shizuka and Johnson 2020). Indeed, empirical links between social position and survival or mortality in systems such as dolphins, killer whales and feral horses support the view that population change can be socially patterned (Stanton and Mann 2012; Nunez, Adelman, and Rubenstein 2015; Ellis et al. 2017). Socially central individuals can support many local and indirect pathways through a network, and their loss may therefore disrupt local clustering or community structure more strongly than the loss of peripheral individuals. In our analyses, low-to-high removal sometimes increased clustering in the absence of compensation, which likely reflects the loss of weakly connected peripheral nodes and the retention of a more cohesive social core. The importance of removal pattern is consistent with broader network-theory based predictions that the effects of perturbation depend on whether loss is random or socially targeted (Albert, Jeong, and Barabasi 2000; Artime et al. 2024). Previous work within single species has shown similar patterns, with social networks often appearing more robust to random loss than to targeted loss of socially central individuals (Lusseau 2003; Williams and Lusseau 2006), and here we show how these effects are mediated by individual social responses.

Allowing individuals to compensate for node loss by forming new connections changed the effect of removal on network structure. Across all four metrics, adding new ties often increased the structural departure from the pre-removal network relative to removal alone. For clustering and modularity, new edges may connect individuals that were not previously embedded within the same local neighbourhood or community, reducing local closure and compartmentalisation. For degree heterogeneity and assortativity, compensation tended to reduce differences among nodes in the number of associations relative to the mean, while lower assortativity indicated a weaker tendency for connected individuals to have similar degrees.

Compensation in response to individual loss could mitigate network changes differently depending on the compensation mechanism, yet we found that compensation had similar effects regardless of mechanism. Friend-of-friend compensation often had less extreme effects than random or degree-biased compensation, which may reflect its bias towards pairs with shared associates (potentially through an ability to maintain the spatial or grouping structure of the network). Regardless, differences among compensation rules were smaller than differences between compensation and no compensation. This is a useful finding for predictions of the effects of individual loss on populations, i.e. for these unweighted global metrics, the act of replacing lost social opportunities by adding new edges had more influence than the exact rule used to place those edges. Going forward, a useful next step may be to test whether these structural similarities also extend directly to patterns of information and disease spread by modelling the spread of simple and complex contagions, as this may then help determine how the consequences of individual loss can be predicted even without detailed knowledge of how surviving animals reconnect.

Further research in real-world networks may illuminate the range of real-world compensation strategies, the ways in which compensation mediates the effects of node loss, and the biological implications of such changes. The complexity of network processes means these implications are likely to depend on context. For instance, lower modularity could promote integration and information flow across the network but could also reduce the compartmental structure that can limit epidemic spread (Sah et al 2018; Evans et al. 2021; Wijayatilake et al. 2026). As such, compensation should be interpreted as a potential mechanism of social reorganisation, with consequences that may be beneficial, costly or neutral depending on the process being considered.

Interpreting compensation as reorganisation aligns with empirical evidence that animals can adjust their social behaviour after losing associates. In wild great tits, individuals increased associations with other flockmates after the loss of social partners (Firth et al. 2017, Abraham, Sheldon, and Firth 2025); in wild house mice, social structure showed community-level resilience after a natural turnover event (Evans et al. 2020); in redfronted lemurs, social disruption was followed by changes in social behaviour (Pfaff et al. 2023) and geese in culled groups increased association strength and formed compensatory social ties (Downing et al. 2023). Social responses to loss have also been described in human social networks, where bereavement can be followed by connective recovery through other relationships (Hobbs and Burke 2017). As such, across systems, compensation may involve forming new ties, strengthening existing ties or reallocating social effort among remaining partners.

The network changes induced by removal and compensation mostly remained significant after modelling accounted for phylogenetic relationships between network species. Phylogenetic structure otherwise explained a large proportion of remaining variation in our models, likely because networks from more closely related species may be more similar in structure. The robustness of our observed effects to the phylogenetic pseudoreplication within our data suggests that understanding the possible effects of compensation within wild networks has cross-context relevance, and that similar patterns may occur within a number of different taxa. However, the significance of metric start value in three of the metrics, in combination with the high amount of variation explained by phylogeny, suggests that more similar networks and species are still likely to express more similar responses to removal and compensation. Comparisons of multiple populations within and across species, using consistent network sampling methods, could now be useful in identifying which features of social organisation predict responses to individual loss, and whether these relationships extend to distantly related species with similar network structures.

Real social responses are likely to be more complex, and individuals may compensate by strengthening existing relationships, reallocating time among remaining partners, forming new ties only after delays, or changing different types of interaction in different ways (Firth et al. 2017; Evans et al. 2020, Farine 2021, Annagiri et al., 2017, Pfaff et al. 2023, Newman et al. 2026). As such, they may increase the strength of existing connections, increasing investment rather than forming new associations. An individual may maintain their “social budget” i.e. total investment across social relationships before and after individual removal, yet this has yet to be comprehensively tested. Future work on weighted, directed and multilayer networks could test whether animals maintain something like a social budget after loss, and whether alternative allocations of social effort have different consequences for transmission, cooperation or information flow.

The amount and cause of loss may also influence whether compensation is feasible, and individual behavioural responses to the loss of associates may depend on the natural turnover of individuals in a population through birth, death or migration (preprint Madsen 2025). At low levels of individual loss, surviving animals may have enough social options to replace some lost associations. At higher levels, there may be too few remaining partners or too few feasible dyads to recover previous levels of connectivity, as reflected by the inability of some simulated networks to accommodate the prescribed number of new edges at high removal percentages. Experimental and observational studies of social perturbation suggest that animal societies can sometimes absorb substantial disturbance, but social instability can also have group-level costs or alter social organisation once disturbance becomes large enough (Flack et al. 2006; Barrett, Henzi, and Lusseau 2012; Franz, Altmann, and Alberts 2015; Maldonado-Chaparro et al. 2018). Disruption of social behaviour may also contribute to Allee effects in social species, where reductions in population size or density reduce individual fitness through the loss of social benefits (Angulo et al. 2018). In further work, compensation could be considered as a capacity that may depend on the scale and speed of demographic change, as well as network state at the time of the loss, rather than as a fixed property of a species or network.

The network state at time of removal, size and scale of loss have been also explored in ecological species networks (where nodes are species and edges interactions between them), where rewiring and adaptive foraging can alter the consequences of species loss, sometimes supporting persistence and sometimes amplifying disruption (Olesen et al 2011; Gilljam, et al 2015; Valdovinos et al. 2016; Su et al 2024). Indeed, in ecological interaction networks, robustness to species loss often depends on network architecture, secondary extinctions and the potential for interaction rewiring (Bascompte and Stouffer 2009; Gilarranz et al. 2017; Morton et al. 2022). Comparisons of animal social networks with ecological networks may be a useful parallel for highlighting general features of networked ecological systems where perturbations can change interactions among the units that remain, and whereby surviving individuals can actively change their own social connections to compensate. In animal social networks, the nodes are individuals within a population that can respond behaviourally to the loss of neighbours, and social structure is therefore not only a property that is affected by demographic decline, but also a possible pathway through which individuals respond to decline. This perspective may be important in considering why compensation may change the structural consequences of loss without necessarily restoring the original network state, and how social behaviour should be considered alongside demographic and ecological processes when predicting the response of animal populations to disturbance.

The impacts of this research are important for population management and prediction of population declines. Management actions that remove individuals, such as culling, translocation, targeted disease control or selective harvesting, may alter network structure through both the removals themselves and the responses of survivors. The two centrality-based methods used to remove individuals in this study show that removing central individuals decreases clustering whilst the removal of peripheral individuals leads to increase in clustering coefficient when compensation does not occur. Population impacts may be species-dependent, and may change the effectiveness of these population management strategies. Social network approaches are increasingly used to identify individuals that may be important for pathogen transmission or information flow, but interventions based on social position could have unintended consequences if remaining individuals reconnect after removal (Silk et al. 2017; Sah, Mann, and Bansal 2018). For example, research on wildlife disease illustrates why social and demographic processes need to be considered when assessing individual removal as a control strategy (tuberculosis in badgers (Weber et al. 2013), infectious facial tumours in Tasmanian devils (Hamede et al. 2009, Drawert et al. 2022), rabies in bats (Streicker et al. 2012)). Our findings may provide a foundation for further work in reducing the uncertainty in the impact of individual removal for disease management but species-specific social traits must be considered, and we suggest that predicting the outcome of network-based interventions may therefore require models that include post-removal social reorganisation, especially when removals are biased towards central or peripheral individuals.

Declining populations lose individuals, but they may also lose social relationships, information pathways, cooperative structure and behavioural traditions (Brakes et al. 2019, Jarić et al. 2026). Social network theory has been proposed as a useful tool for conservation because population viability can depend on social processes as well as abundance (Snijders et al. 2017). Our analyses suggest that compensatory social behaviour may be one mechanism through which populations respond to such losses, although compensation may generate new network states with different functional consequences with potential adaptive benefits. Furthermore, our results show that we may not need to know the precise compensation strategy in order to predict the response in a social network. Individual loss in animal societies can be considered as a dynamic social perturbation, whereby demographic change removes individuals and ties, and surviving animals may then reshape the remaining network through their own behaviour. Accounting for compensatory social behaviour may improve predictions of how animal societies respond to demographic change in increasingly disturbed populations. As such, these findings highlight the value of incorporating social responses into predictions for threatened populations, particularly in an era of mass extinction (Morton et al. 2022), although social responses are often absent from studies of population decline (Spooner et al. 2018, Cowlishaw et al. 2008, Collen et al. 2011).

## Methods

### Network selection

Real world wild animal social networks were accessed from the Animal Social Network Repository (Sah et al. 2019, Collier et al. 2024 [Accessed 06/06/2025]). Only networks with greater than 10 nodes were retained. We also removed networks with greater than 50% isolated nodes, networks with edges based on spatial proximity, and networks from captive populations. After this filtering, 268 networks were retained for analysis (Supplementary Information Figure SI.1). Throughout the analysis, all networks were treated as unweighted and undirected in order to simplify simulation.

### Removal

Each removal represents the loss of an individual from a social network - in a wild system, this could represent death, removal or emigration. Three methods of removal were simulated: random, ‘high-to-low’ and ‘low-to-high’ centrality. Random removal provided a form of null comparison for the centrality-based removals described below, with fewer assumptions. Removal was also simulated to target nodes based on their centrality in the original network, with one method targeting nodes in order of decreasing centrality (high-to-low centrality) and the other in increasing centrality (low-to-high centrality). Eigenvector centrality was used as the centrality metric, calculated using the igraph function (Antonov et al. 2026 [Version 1.3.5]).

To investigate how the effects of removal and rewiring might vary with the extent of node loss, we simulated three levels of node removal: 10, 20 or 30% of the original network size (Figure 1B). The nodes lost at each level were calculated independently, rather than sequentially i.e. the 20% nodes removed were an independent selection from the original full set, rather than the set of nodes removed in the 10% simulation plus an additional 10%. With 3 removal levels across each of three removal methods, each network generated 9 reduced networks with nodes removed. As random node removal could lead to stochastic explanations for our results, we accounted for this by repeating each removal, compensation and analysis process three times, with three different random seeds (we were constrained from further randomisations by computational limitations). All results and analyses presented in the main text were conducted with random seed 1, as there was little variation in results between seeds (provided in the Supplementary Information).

### Compensation strategies

We simulated three compensation strategies to reduce the impact of our assumptions on the results, and to allow us to compare the importance of the compensation method for the final structure of the reduced network. Each of the three methods followed the same basic rules. The number of edges to be added is determined by the number of edges lost by the remaining individuals (Figure 1C). A probability matrix was calculated to determine where the edges were added, with no edges allowed between individuals which were already associated or from an individual to itself. Each compensation method had a different calculation method, where the probability of an edge being added between a pair of nodes was weighted according to a specific compensation strategy and was simulated 100 times for each network (Figure 1D).

‘Random’ compensation added new edges at random between individuals which were not already associated. ‘Degree’ compensation was randomly generated, weighted by the number of edges lost for each remaining node-pair. This meant that edges were more likely to form between nodes which were further from their original degree (number of associates). ‘Friend of friend’ compensation was similarly weighted by lost edges for each node-pair but further multiplied based on the number of neighbours the node-pair had in common. These probability-based rules do not require individuals to recover their original degree. We also calculated a control of no compensation, with no new edges added after removal.

At a certain point, our compensation algorithm becomes impossible when the number of edges to add exceeds the number of possible connections whilst avoiding multiple connections and self-loops. In this case, compensation is considered infeasible or “collapsed” under the specified rules and network metrics are treated as missing values.

Simulations were originally conducted from 0% to 90% removal. However, the exclusion of networks for which compensation was infeasible at higher removal percentages (see Supplementary Information) meant that sample sizes were much lower after 30% node removal (below 200 in random removal). We therefore restricted the main analyses to 0-30% removal to limit the loss of networks from the comparison. Descriptions and analyses of all results on the complete (0-90% removal) dataset are provided in the Supplementary Information.

### Measuring responses

To compare network structure before and after node removal, we selected four global network metrics (Figure 1E). As removal inherently changes network structure through a reduction in node number, we aimed to select metrics which were more robust to network size, both in node number and edge number. The final metrics used for the analysis were clustering, modularity, degree assortativity and degree heterogeneity. Definitions and real-world relevance for each metric are provided in Table 1.

**Table 1:** Definitions, real world effects, and calculation methods for each of the metrics used within this analysis.

| **Metric** | **Definition** | **Effect on social processes** | **Effect on vulnerability to removals** | **R calculation function** |
|---|---|---|---|---|
| **Clustering** | The tendency of nodes to form local clusters (averaged across the network). | Networks may reduce clustering coefficient in response to disease (Croft et al. 2011). Clustering is likely to increase transmission of disease or information within local groups, but reduce spread across the whole network (Wey et al. 2008). | Diverging opinions on whether clustering increases or decreases network robustness (Artime 2024). Real world networks may, after removals, maintain stable clustering (Lusseau 2003) or increase clustering (Manno 2008). | ‘transitivity’ from igraph (Csardi and Nepusz 2006 [Version 1.3.5]), with type set as local average, then averaged across the network. |
| **Modularity** | The extent to which a network can be partitioned into distinct communities. | Modular structuring represents important social information, and has implications for the spread of disease and information (Evans et al. 2021), and other social processes such as cooperation (Marcoux and Lusseau 2013). | Modularity may increase the vulnerability of a network to removals (Kim and Cho 2015; Nguyen et al. 2021); however, other research has shown that modularity may ‘contain’ permutations (Gilarranz 2017). Work in ecological networks has shown rewiring to reinforce modular structure (Su, Ma, and Hui 2024). | ‘modularity’ from igraph (Csardi and Nepusz 2006 [Version 1.3.5]). |
| **Degree heterogeneity** | The variation in number of associations between individuals in the network. | High values of degree heterogeneity are associated with smaller and shorter epidemics (Sah, Mann and Bansal 2017) | Very few papers measure degree heterogeneity, yet Franz et al. 2015 found it recovered and became stable. | sd(degree)/mean (degree) where degree is the number of associates for each node. |
| **Degree assortativity** | The tendency of nodes with a large number of connections to connect with nodes that also have large numbers of connections. Number of connections (degree) is normalised across the network. | Both information and disease may spread more quickly through an assortative network (Noldus and Van Mieghem 2014). | Analysis has shown that networks with a higher amount of degree assortativity are more robust to node removals (Newman 2003). | ‘assortativity_degree’ from igraph (Csardi and Nepusz 2006 [Version 1.3.5]), normalised between 0 and 1 |

### Analysis of results

Simulations allow us to assess patterns within the ‘world’ of the simulation, and cannot necessarily demonstrate biological effects, so simulation results should be explored with care (see Evans, Heppenstall and Birkin (2017) and White et al. (2013) for discussion). In this context, statistical ‘significance’ could be generated artificially by increasing the number of simulations performed. We therefore present results in a number of ways to draw out conclusions from our simulated data. First, we present changes in network metrics as a percentage change in the mean value between compensation methods, removal methods, or removal percentages (for example, percentage change in mean clustering coefficient from 0% to 30% removal). We then use permutation tests to assess these observed changes, and regression models with phylogenetic random effects to further investigate the role of pseudoreplication in our results.

### Percentage change

As discussed above, changes in network metrics are presented as percentage change in mean values. Additionally, we provide simulation intervals in the figures, which estimate the range around the mean metric value. Following Cheng and Holland (2004), we calculated simulation intervals under the assumption that the only relevant source of variability in our model was *simulation variability*, generated from the random selection of removal nodes. We calculated simulation intervals around the mean metric values as the 2.5% quantile to the 97.5% quantile, to estimate 95% of the variation. These intervals ignore the variation within networks, as each network metric value is itself a mean.

### Permutation tests

To help assess whether observed differences in simulation outputs were meaningful, we conducted permutation tests of the percentage differences between mean values, using a protocol which is described in the Supplementary Information, along with the complete results from this analysis. Where relevant, these values are reported in the text as *p_perm_* where n.s denotes a p_perm_ greater than 0.05. A significant p_perm_ represents a percentage change significantly greater or smaller than would be expected if removal method, rewiring method, or removal percentage were randomised. This permutation protocol was used to avoid the issues with using traditional significance to assess simulation results, and provides an indication of when a result should be considered meaningful, supported by visual assessment of the magnitude of the difference between results (White et al. 2013).

### Accounting for phylogeny

We then used random effects to analyse the contribution of network pseudoreplication to the variation in network responses to removal and compensation. This was in order to explore the sources of variation between networks. Specifically, we analysed log multiplication factor (MF), where MF represented the scale of change in mean metric value between 0% and 30% removal, calculated as follows:

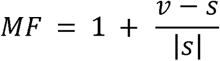

where *v* is the network metric value at 30% removal and *s* is the network metric value at 0% removal. The absolute value of s was used to account for network metrics that can take negative values. An MF of 1 therefore indicates no change in metric value from 0% to 30% node removal, whereas values above or below 1 indicate an increase or decrease respectively. We used mean metric value to account for the lack of variation in metric values for networks under high-to-low or low-to-high removal and no compensation. We then log transformed MF in order to avoid a higher weighting of values which were above 1. MF could not be calculated for networks which had an initial metric value of 0, and log MF could not be calculated where MF was zero or negative. Therefore, these networks were not included in these specific models (this removed the following percentage of observations: *clustering* 10.6%; *modularity* 11.7%; *degree heterogeneity* 3.8%; *degree assortativity* 5.0%). As this is possibly an important feature of these networks, these were subsequently investigated further in the Supplementary Information.

We aimed to investigate the extent to which variation across networks in the effect of node loss and compensation were associated with three potential sources of pseudoreplication – network identity, species identity, and phylogenetic relatedness. Our data did not allow these sources of variation to be estimated simultaneously, so we fitted a model for each network metrics with either network ID, species ID, or phylogeny as a random effect. Phylogeny was included in the model as a covariance matrix calculated from a phylogeny of the species of the networks used (Hadfield and Nakagawa 2010), estimated with ‘rotl’ (Michonneau, Brown and Winter 2016) and ape (Paradis and Schliep 2019) (one network from the species *Acutisoma proximum* was removed for these models as this species was not available in the Open Tree of Life).

To account for variation generated by simulation processes, we included an interaction between removal method and compensation method, and the start value of the metric at 0% removal, as fixed effects. We used brms (Bürkner 2021) to run these models. All model families were Student’s t, with normal (0,1) priors for fixed effects and default brms priors otherwise.

This analysis found that for all four metrics, phylogeny was the random effect explaining most variation after accounting for fixed effects (over 75% of residual variation in all four metrics, see Supplementary Information). We were interested in whether the results we obtained from permutation testing were upheld by models which accounted for phylogenetic pseudoreplication, so in the main text we present the fixed effect estimates of log multiplication factor at 30% removal, for each four metrics.

## Acknowledgements

We would like to acknowledge the team behind the ASNR and all the researchers who collected and shared the original data included in the ASNR.

## Data and code availability

Data and code is available at Figshare with doi.org/10.6084/m9.figshare.34995862

