## Supplementary Information for "Social compensation changes predicted effects of individual loss in animal societies"

**Affiliations**:
1. Department of Biology, Life and Mind Building, University of Oxford, Oxford, OX1 3JS
2. School of Biology, University of Leeds, Leeds, LS2 9JT

#### Network selection

Figure SI.1 shows the numbers of networks from ASNR filtered out according to various selection processes, in order to have 268 networks for analysis. Networks were filtered to be greater than 10 nodes with network density not equal to 1 in order to ensure that removals were always whole numbers, and to make sure that edge compensation would be meaningful within the network. We only analysed wild populations with networks that were not derived from spatial proximity, as we were only interested in these effects on wild populations, and spatial proximity networks didn't reflect the sort of detailed social data we were interested in. We discarded networks with greater than 50% isolated nodes to again ensure that compensation is meaningful, and further removed networks which did not align with the metadata from ASNR, to avoid errors. After this filtering, we retained 268 networks which were analysed in this report.

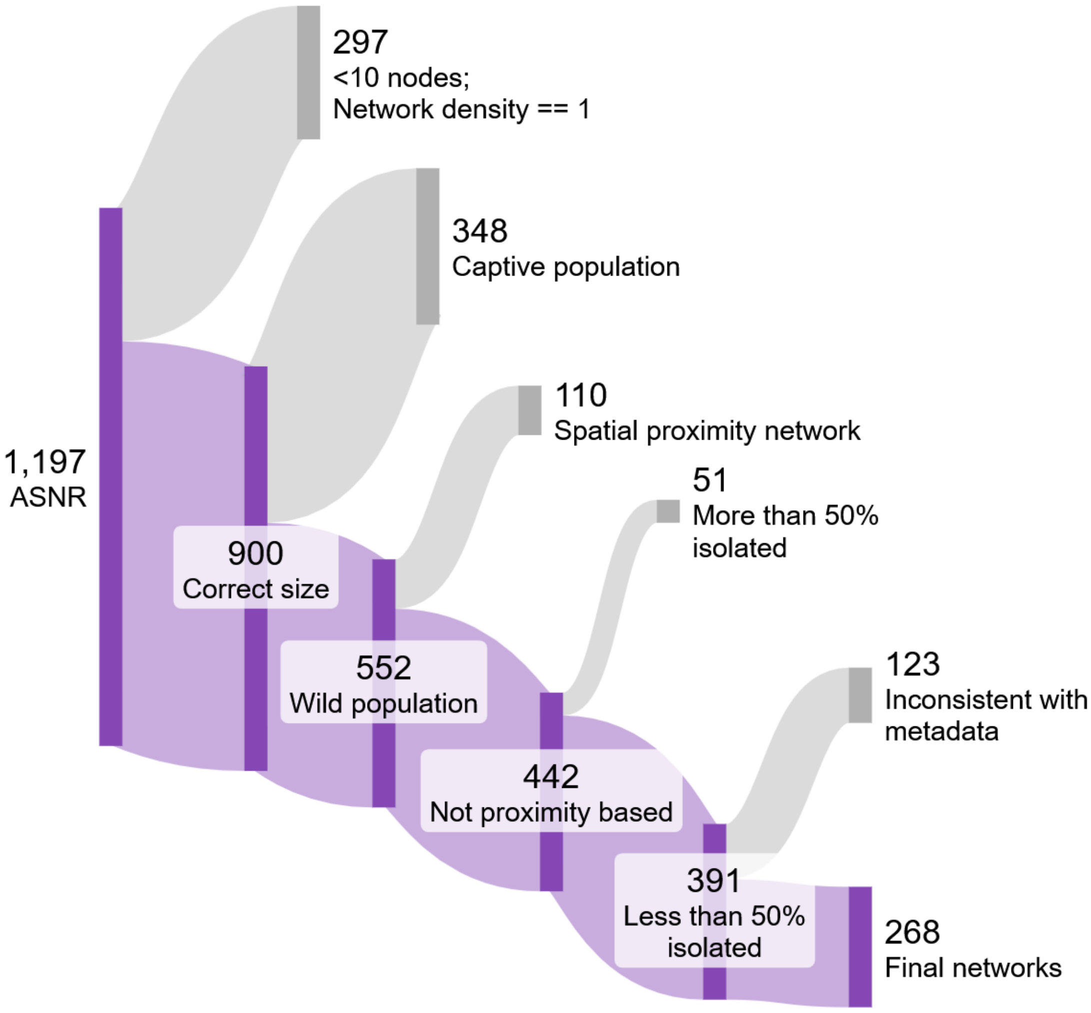

*Figure SI.1: A diagram of the number of networks retained for analysis after various filters were applied, with retained networks in purple and discarded networks in grey.*

*Table SI.1: Network sample size at different node removal percentages, removal methods, and compensation methods. 'Sample size' refers to the number of networks which were able to compensate, and were therefore included in our visualisations and analyses. Numbers here refer to random seed 1, the results of which are presented in the analysis.*

| **Removal method** | **Compensation method** | **Sample size at removal %** | | | |
|---|---|---|---|---|---|
| | | 0 | 10 | 20 | 30 |
| Random | Random | 268 | 250 | 252 | 238 |
| | Degree | 268 | 253 | 254 | 238 |
| | Friend of friend | 268 | 253 | 254 | 238 |
| | No compensation | 268 | 268 | 268 | 268 |
| High-to-low | Random | 268 | 249 | 254 | 254 |
| | Degree | 268 | 249 | 254 | 254 |
| | Friend of friend | 268 | 249 | 254 | 254 |
| | No compensation | 268 | 254 | 254 | 254 |
| Low-to-high | Random | 268 | 238 | 238 | 246 |
| | Degree | 268 | 241 | 241 | 249 |
| | Friend of friend | 268 | 242 | 241 | 249 |
| | No compensation | 268 | 254 | 254 | 254 |

#### Comparison between random seeds

In order to assess whether the random element in our analysis impacted our results, we ran the analysis with two additional random seeds. We then visually compared results, and determined that any differences due to randomness were negligible. For ease of interpretation, we provide only the results from random seed 1 in the main text, but provide the main text figures made with random seeds 2 and 3 here. Differences are expected only under random removal, as removal from high-to-low and low-to-high centrality did not have a stochastic element.

*Seed 2*

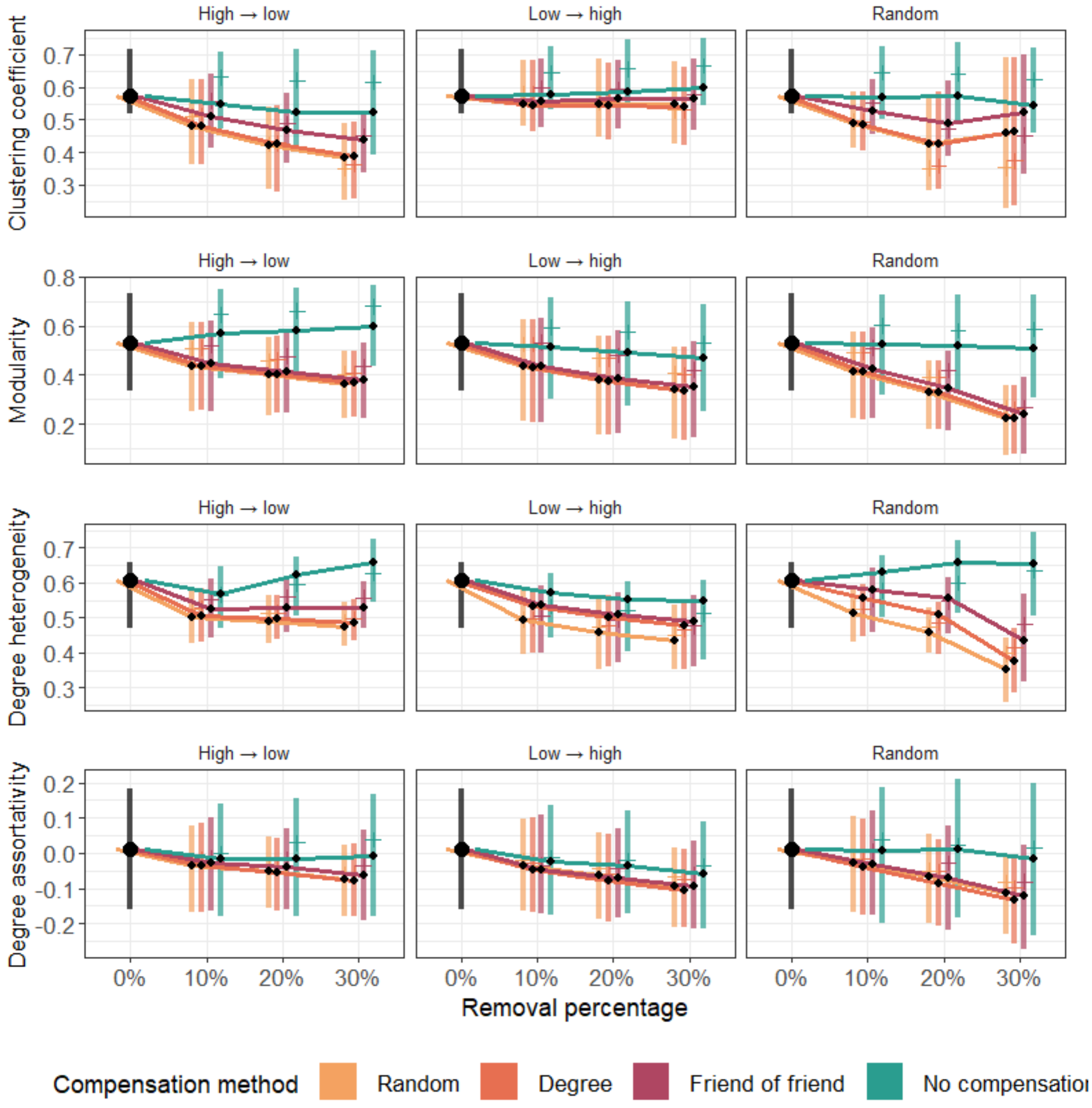

*Figure SI.2: Comparison of network metrics under random seed 2 for three removal methods; high-low centrality, low-high centrality and random removal. Compensation method is shown with no compensation (teal), random compensation (yellow), preserving degree (orange), and bias to friends of friends (red). As all networks started at the same metric values, metrics at step 0 are coloured black. Lines between black points represent mean*

values. Line ranges contain 50% of the data, between the 25% and 75% quantile. Small side bars represent median values.

*Seed 3*

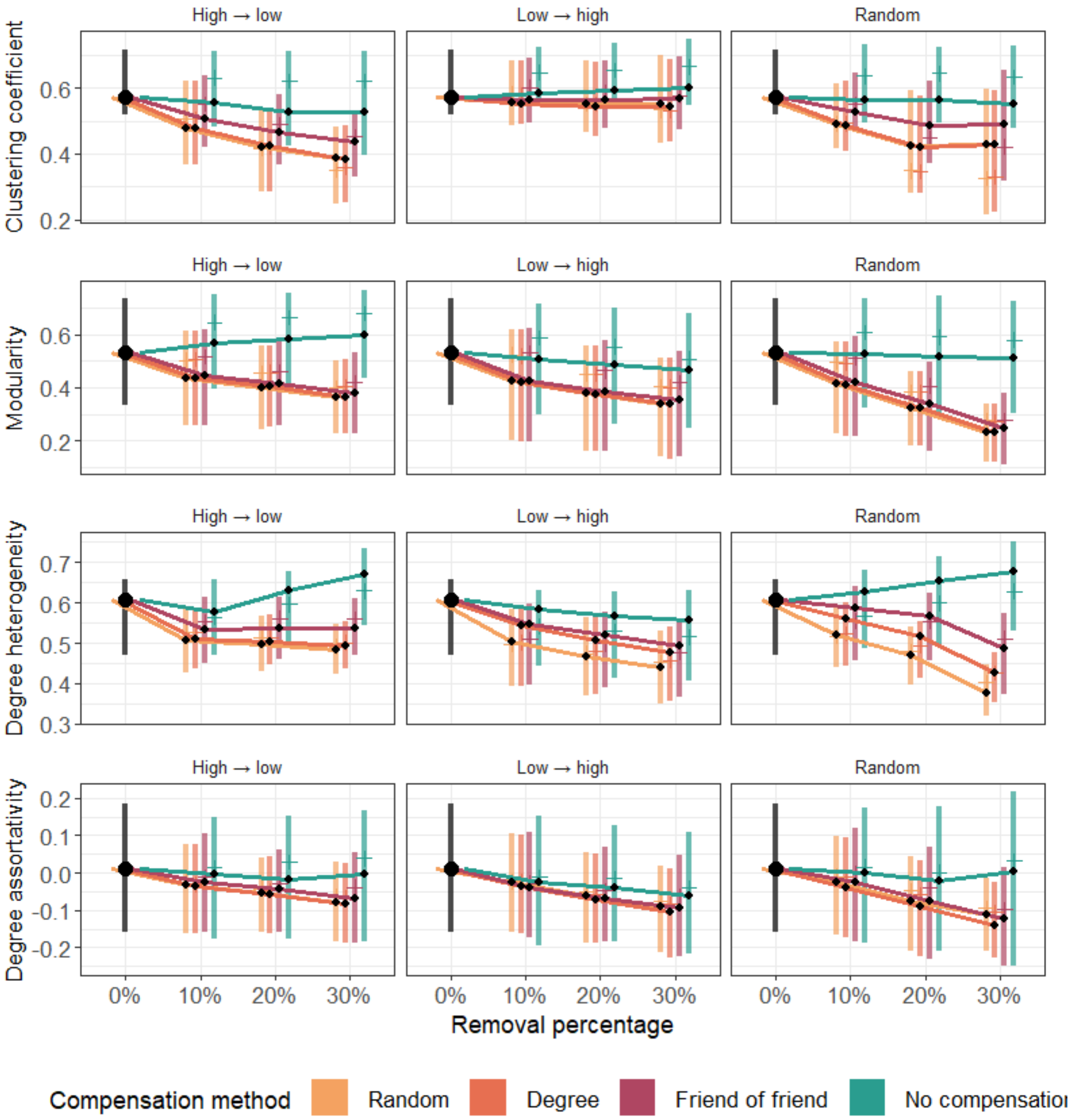

*Figure SI.3: Comparison of network metrics under random seed 3 for three removal methods; high-low centrality, low-high centrality and random removal. Compensation method is shown with no compensation (teal), random compensation (yellow), preserving degree (orange), and bias to friends of friends (red). As all networks started at the same metric values, metrics at step 0 are coloured black. Lines between black points represent mean values. Line ranges contain 50% of the data, between the 25% and 75% quantile. Small side bars represent median values.*

#### Metric results up to 90% removal

We chose to present in the main text node removal results up to 30%. As discussed in the Methods, this was due to reduced sample size after 30% removal and the relative unlikelihood of 40%+ mortality events. Here, we present metric results for removal percentages up to 90%.

##### Clustering

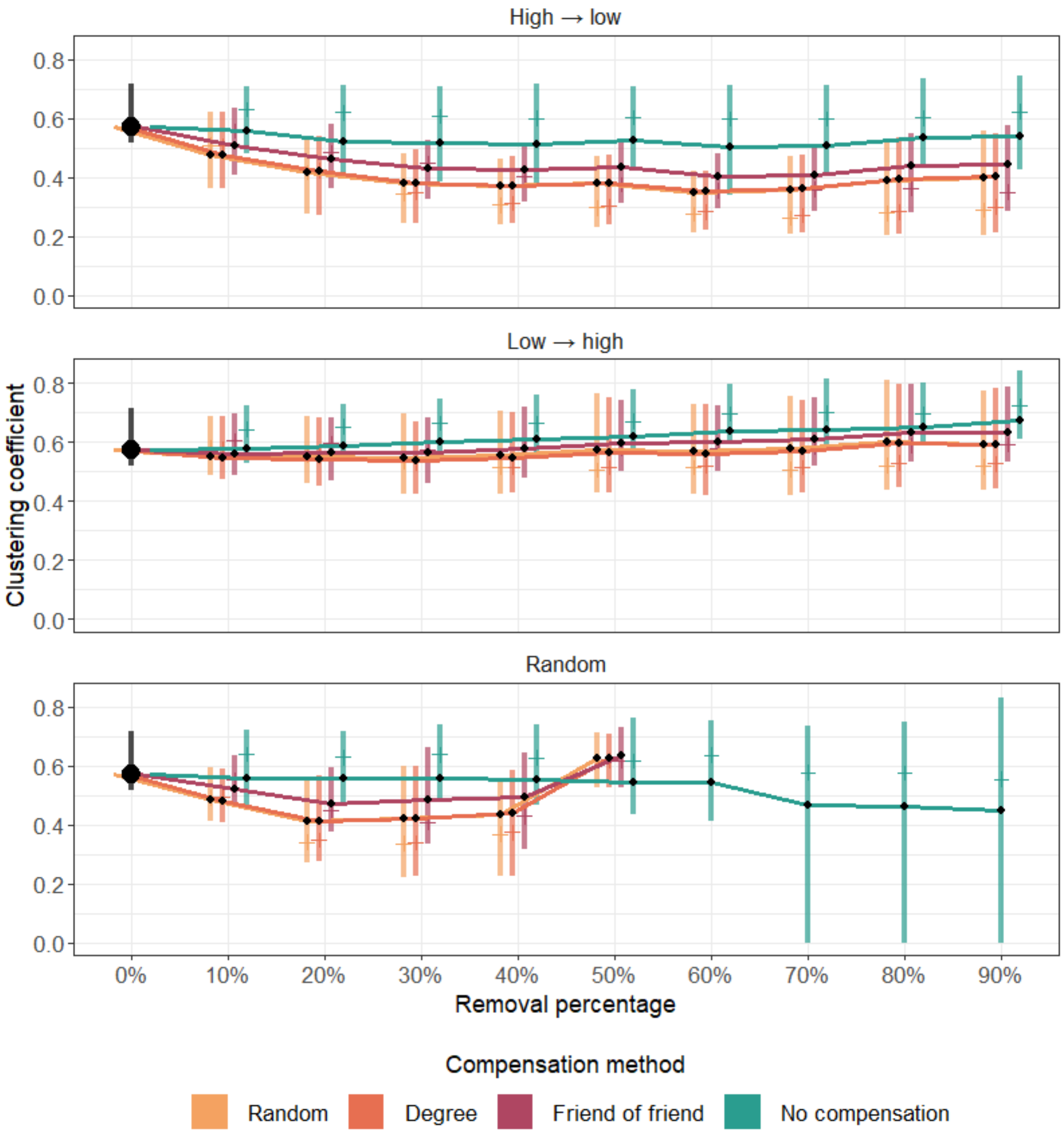

*Figure SI.4: Comparison of clustering up to 90% removal for three removal methods; high-low centrality, low-high centrality and random removal. Compensation method is shown with no compensation (teal), random compensation (yellow), preserving degree (orange), and bias to friends of friends (red). As all networks started at the same metric values, metrics at step 0 are coloured black. Lines between black points represent mean values. Line ranges*

*contain 50% of the data, between the 25% and 75% quantile. Small side bars represent median values.*

#### Modularity

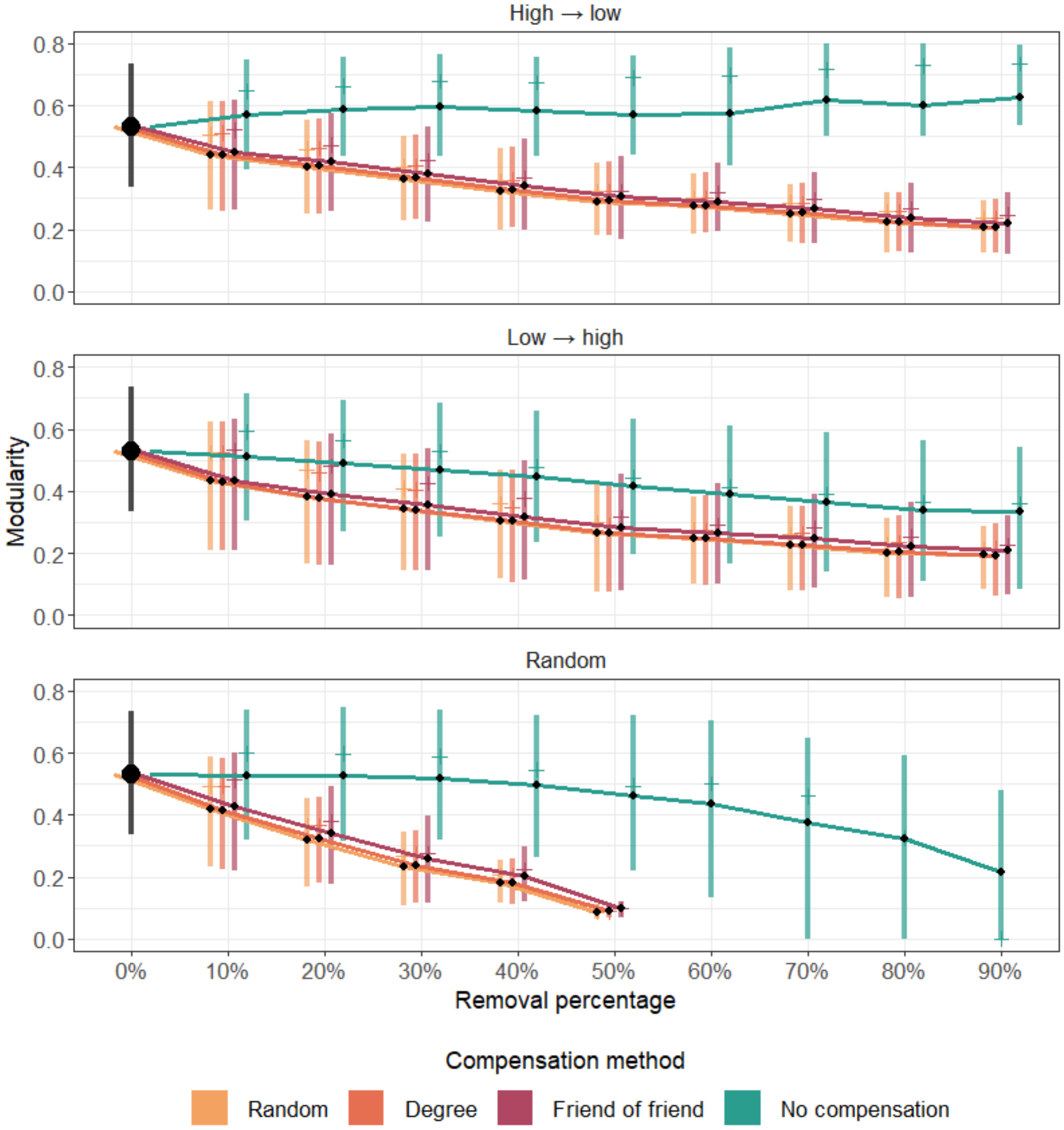

*Figure SI.5: Comparison of modularity up to 90% removal for three removal methods; high-low centrality, low-high centrality and random removal. Compensation method is shown with no compensation (teal), random compensation (yellow), preserving degree (orange), and bias to friends of friends (red). As all networks started at the same metric values, metrics at step 0 are coloured black. Lines between black points represent mean values. Line ranges contain 50% of the data, between the 25% and 75% quantile. Small side bars represent median values.*

#### Degree heterogeneity

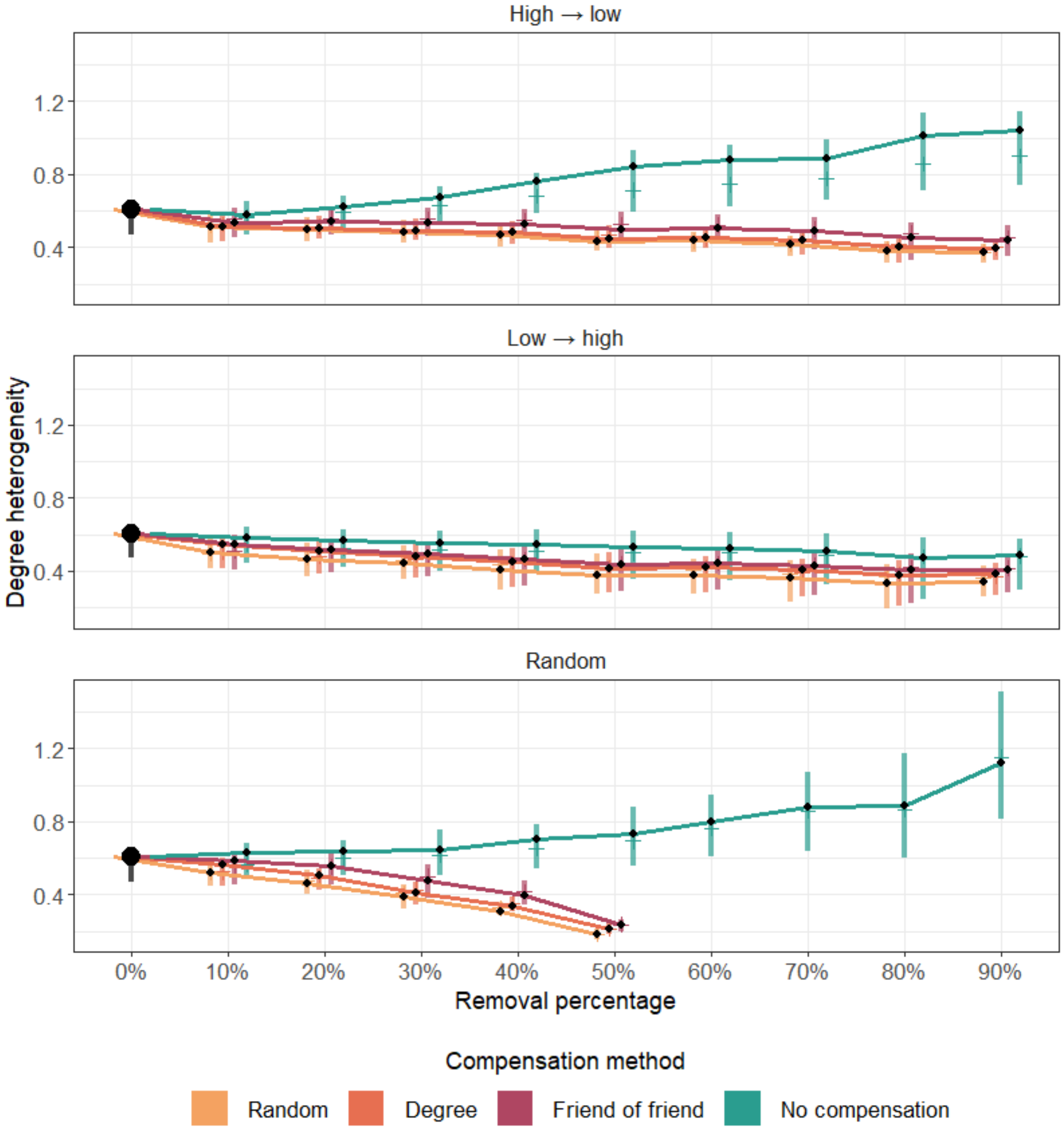

*Figure SI.6: Comparison of degree heterogeneity up to 90% removal for three removal methods; high-low centrality, low-high centrality and random removal. Compensation method is shown with no compensation (teal), random compensation (yellow), preserving degree (orange), and bias to friends of friends (red). As all networks started at the same metric values, metrics at step 0 are coloured black. Lines between black points represent mean values. Line ranges contain 50% of the data, between the 25% and 75% quantile. Small side bars represent median values.*

#### Degree assortativity

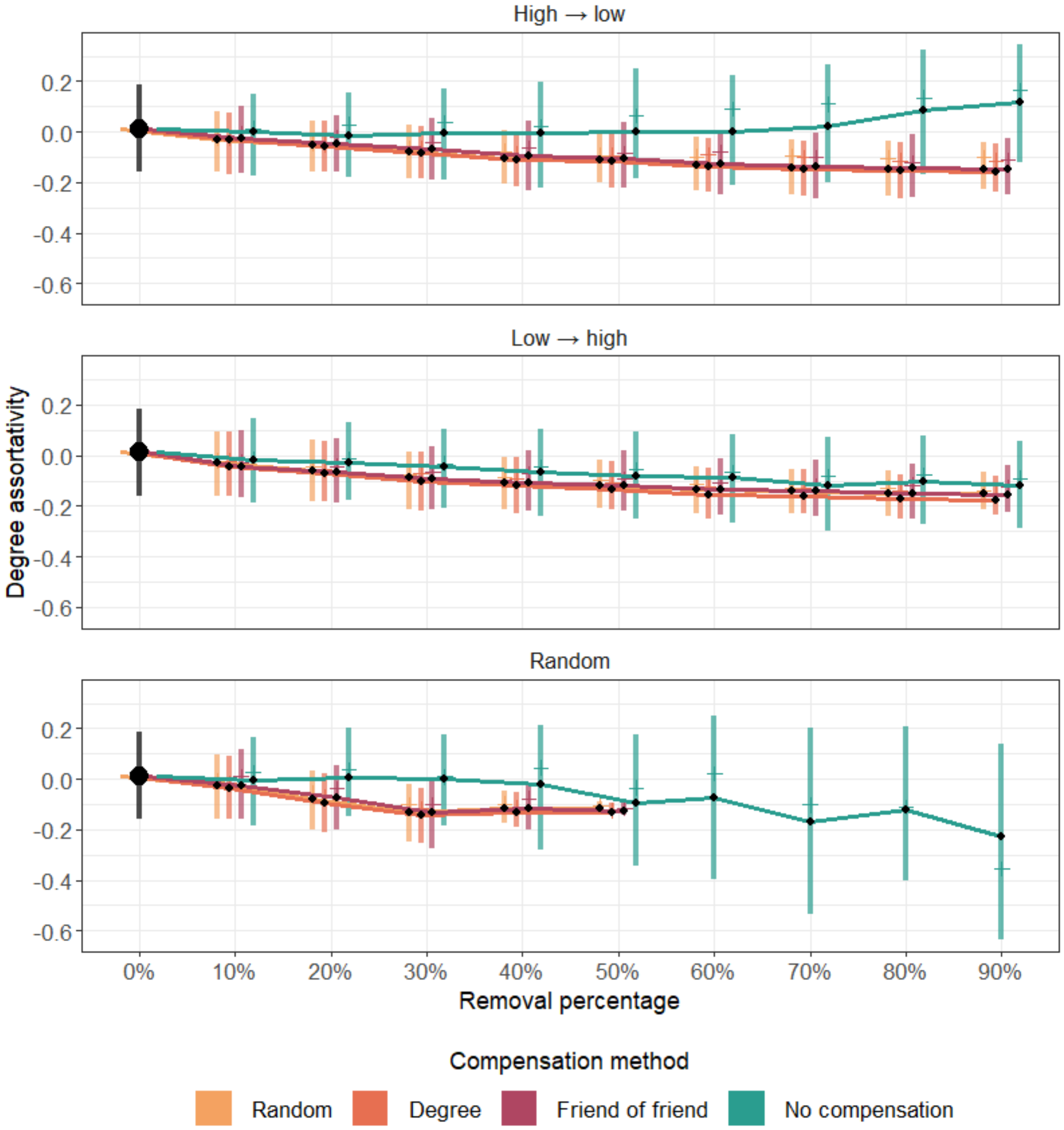

*Figure SI.7: Comparison of degree assortativity up to 90% removal for three removal methods; high-low centrality, low-high centrality and random removal. Compensation method is shown with no compensation (teal), random compensation (yellow), preserving degree (orange), and bias to friends of friends (red). As all networks started at the same metric values, metrics at step 0 are coloured black. Lines between black points represent mean values. Line ranges contain 50% of the data, between the 25% and 75% quantile. Small side bars represent median values.*

### Permutation testing

Permutation tests are a non-parametric test which allow estimates such as p-values to be obtained from data which may not meet the required assumptions for other tests. They work by randomising data across the variables of interest many times. The distribution of results for these randomisations is then compared to the result from the original data to obtain a p-value. A p-value less than 0.05 implies that the observed result was due to the real distribution of variables, rather than random chance or other underlying structures in the data. For our data, permutation tests were used to assess claims about the percentage differences between various categories of data - this tests the significance of the difference itself. The exception to this was for degree assortativity, where we couldn't calculate percentage differences

Our results take three main forms, and a different permutation structure was required to test each of these. The first comparison is (a) comparisons between compensation types within a removal method e.g *"at 30% high-to-low removal, the clustering coefficient is [X]% higher with no compensation than with degree compensation"*. As the factor of interest here is compensation type, permutations to test these conclusions involved randomisation of compensation type between networks, while holding removal method and percentage removal consistent. In other words, within removal methods, one network was given a randomised compensation type across all removal percentages.

The second comparison is (b), comparisons between removal methods within compensation type, e.g *"at 30% removal, clustering coefficient with no compensation in random removal is X% lower than no compensation in low-to-high removal".* Here, removal method was randomised between networks, while compensation type and percentage removal remained stable. Within compensation types, one network was given a randomised removal method across removal percentages.

The third and final comparison, (c), is comparison between removal levels, e.g "*with low-to-high removal, clustering coefficient with no compensation was X% higher at 30% removal than it was at 0% removal"*. For this form of result, removal method and compensation method remained stable while removal percentage was randomised, while keeping network ID consistent. Within removal methods and removal percentages, the step number for each network's metric measures was randomised.

Figure SI.8 provides a guide for reading the permutation analysis tables provided below.

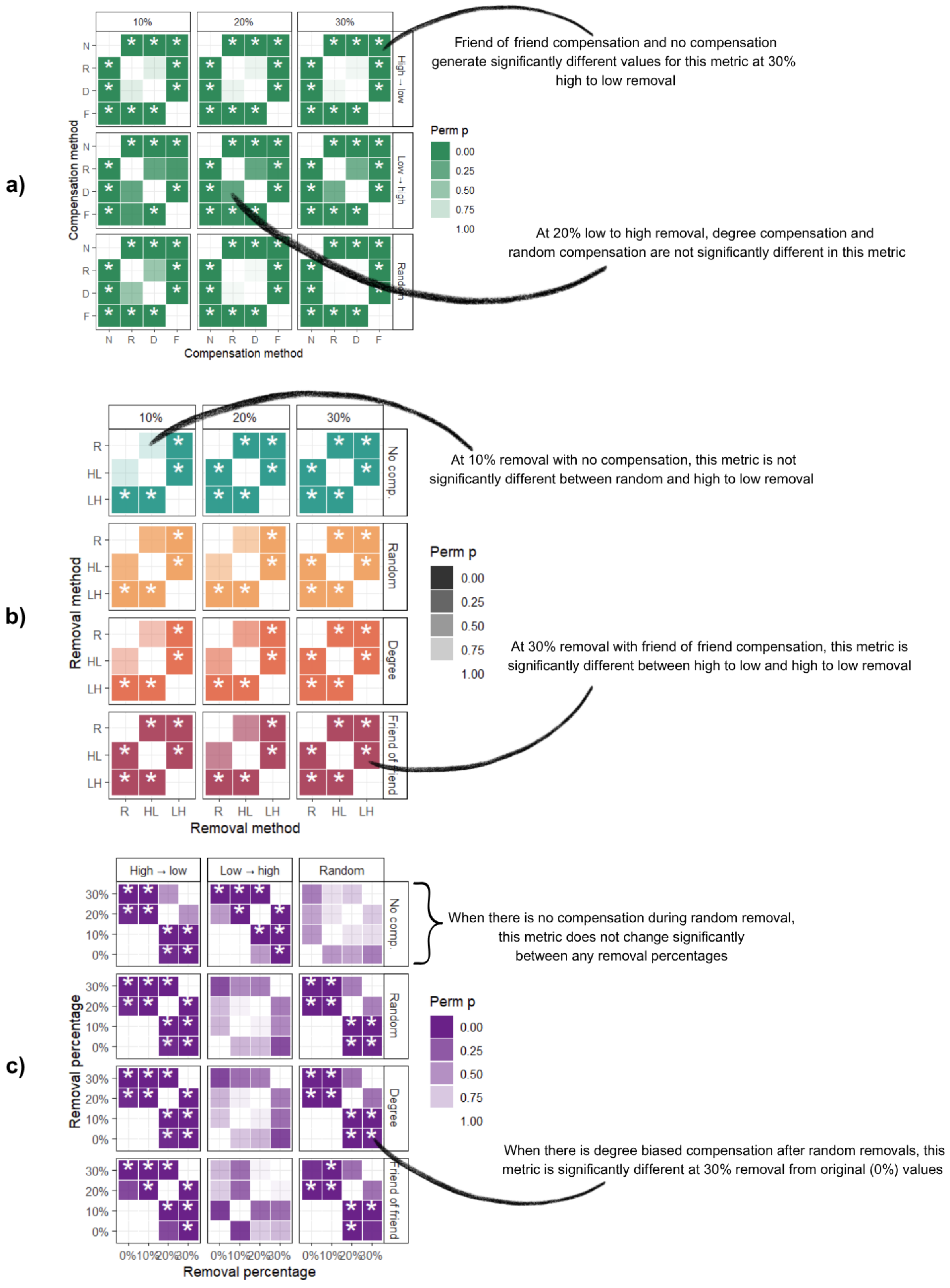

*Figure SI.8: A guide to reading the permutation tables for a) comparison between rewiring methods, b) comparison between removal methods, and c) comparison between removal percentages. Example table cells are labelled with how you would interpret the result.*

***a) Comparison between rewiring methods***

*Clustering*

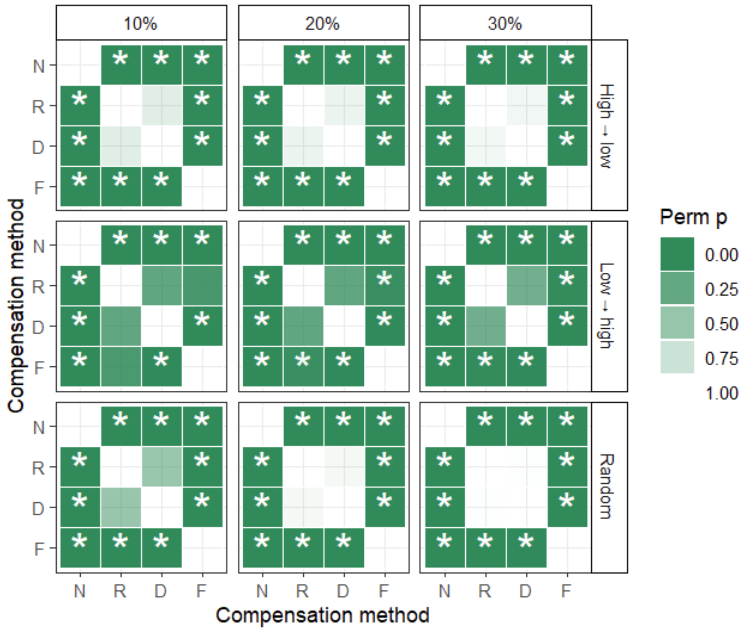

*Figure SI.9: A table of p-values generated by 1000 permutations of compensation type within networks, and calculating resulting percentage changes in clustering coefficient. Stars represent significant ($p < 0.05$) comparisons. Compensation types are N = no compensation, R = random compensation, D = degree biased compensation and F = friend-of-friend compensation.*

*Modularity*

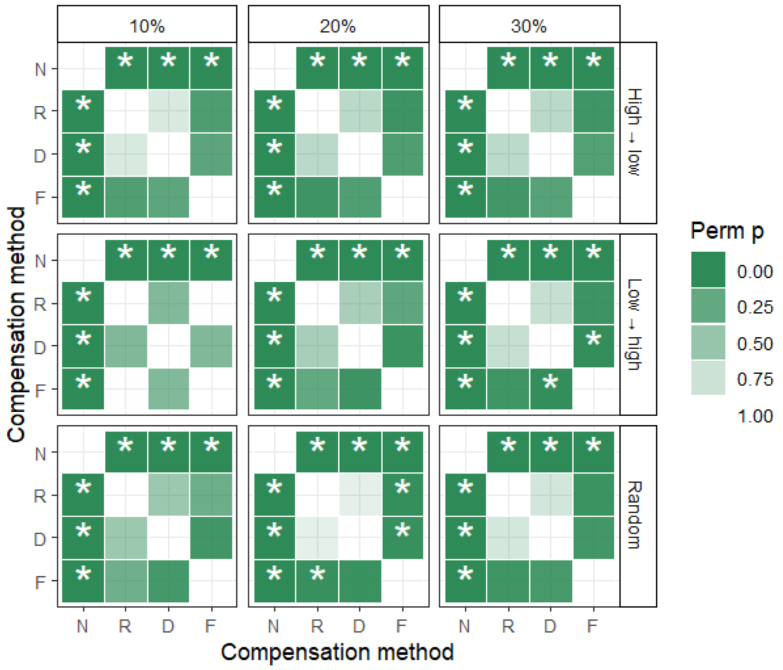

*Figure SI.10: A table of p-values generated by 1000 permutations of compensation type within networks, and calculating resulting percentage changes in modularity. Stars represent significant ($p < 0.05$) comparisons. Compensation types are N = no compensation, R = random compensation, D = degree biased compensation and F = friend-of-friend compensation.*

*Degree heterogeneity*

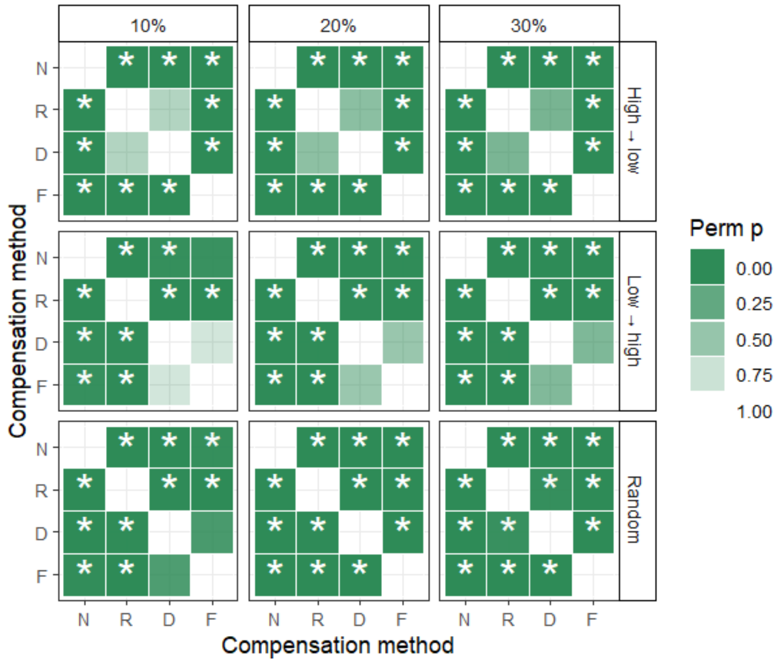

*Figure SI.11: A table of p-values generated by 1000 permutations of compensation type within networks, and calculating resulting percentage changes in degree heterogeneity. Stars represent significant (p < 0.05) comparisons. Compensation types are N = no compensation, R = random compensation, D = degree biased compensation and F = friend-of-friend compensation.*

*Degree assortativity*

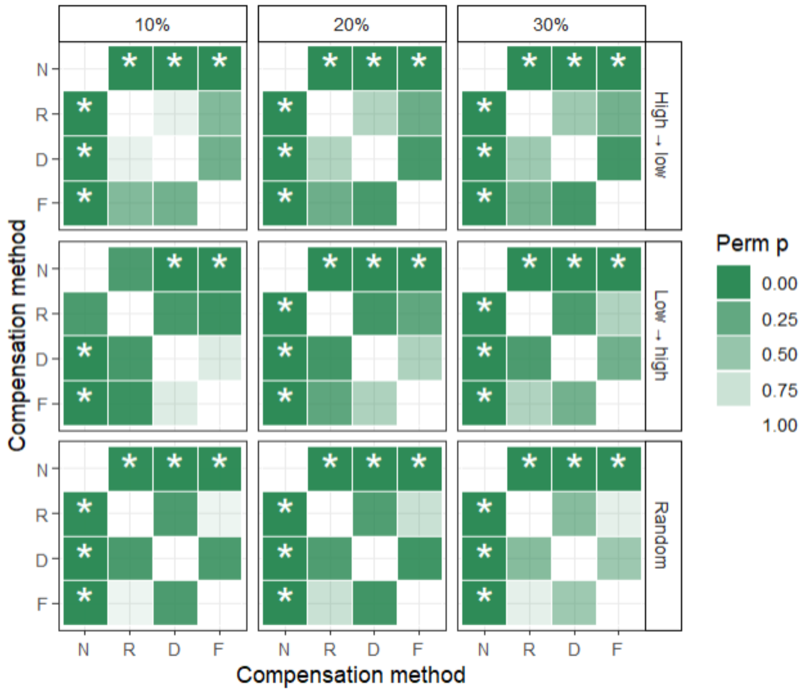

*Figure SI.12: A table of p-values generated by 1000 permutations of compensation type within networks, and calculating resulting percentage changes in normalised degree assortativity. Stars represent significant (p < 0.05) comparisons. Compensation types are N = no compensation, R = random compensation, D = degree biased compensation and F = friend-of-friend compensation.*

***b) Comparison between removal methods***

*Clustering*

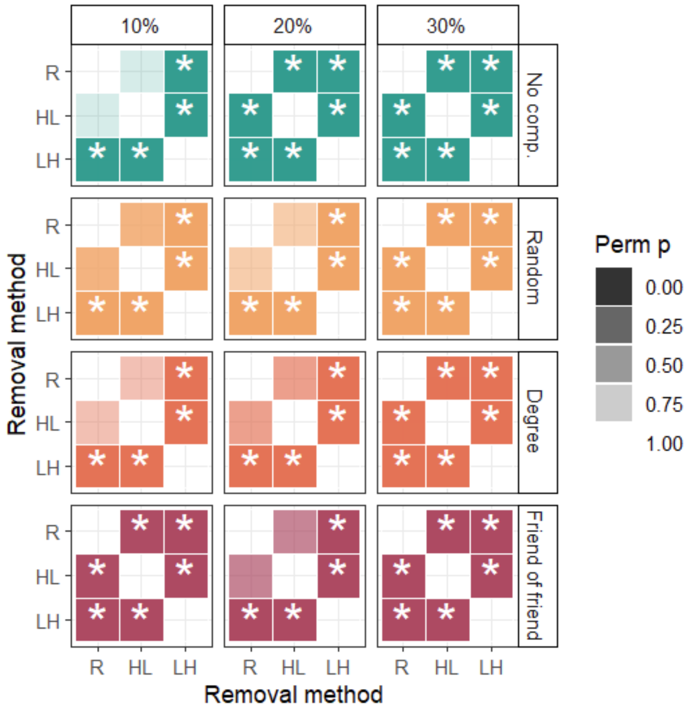

*Figure SI.13: A table of p-values generated by 1000 permutations of compensation type within networks, and calculating resulting percentage changes in clustering coefficient. Stars represent significant ($p < 0.05$) comparisons. Removal methods are R = random; HL = high to low; LH = low to high.*

*Modularity*

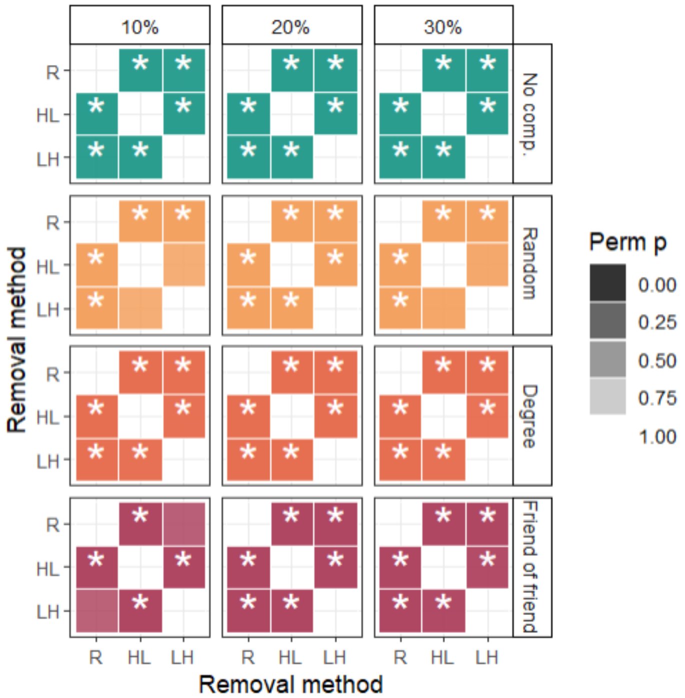

*Figure SI.14: A table of p-values generated by 1000 permutations of compensation type within networks, and calculating resulting percentage changes in modularity. Stars represent significant ($p < 0.05$) comparisons. Removal methods are R = random; HL = high to low; LH = low to high.*

*Degree heterogeneity*

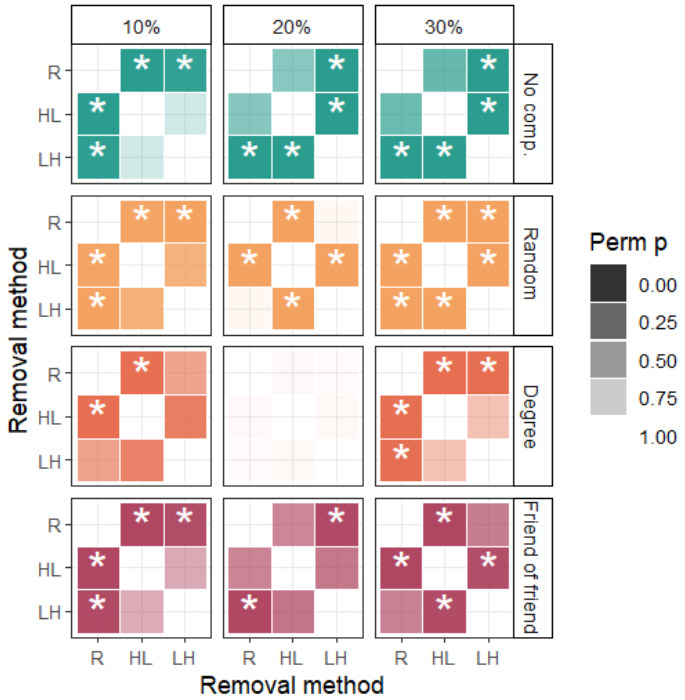

*Figure SI.15: A table of p-values generated by 1000 permutations of compensation type within networks, and calculating resulting percentage changes in degree heterogeneity. Stars represent significant (p < 0.05) comparisons. Removal methods are R = random; HL = high to low; LH = low to high.*

*Degree assortativity*

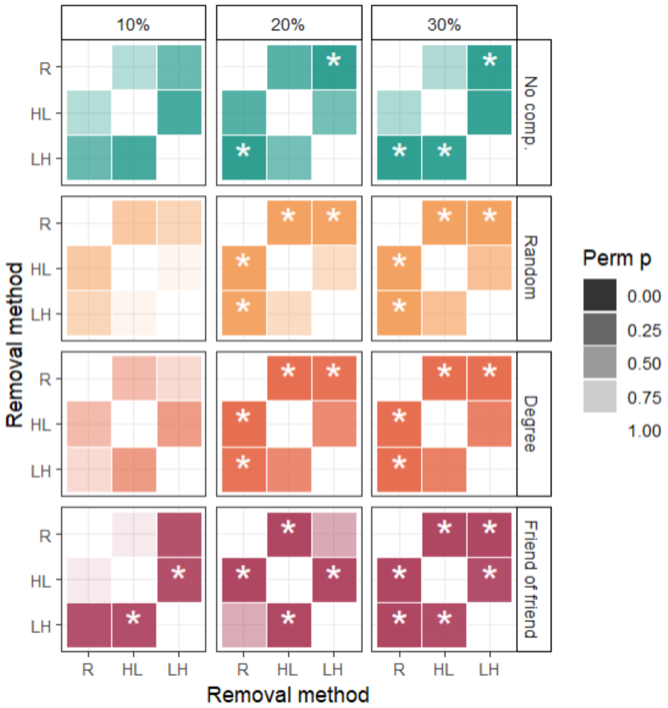

*Figure SI.16: A table of p-values generated by 1000 permutations of compensation type within networks, and calculating resulting percentage changes in degree assortativity. Stars represent significant (p < 0.05) comparisons. Removal methods are R = random; HL = high to low; LH = low to high.*

***c) Comparison between removal percentages***

*Clustering*

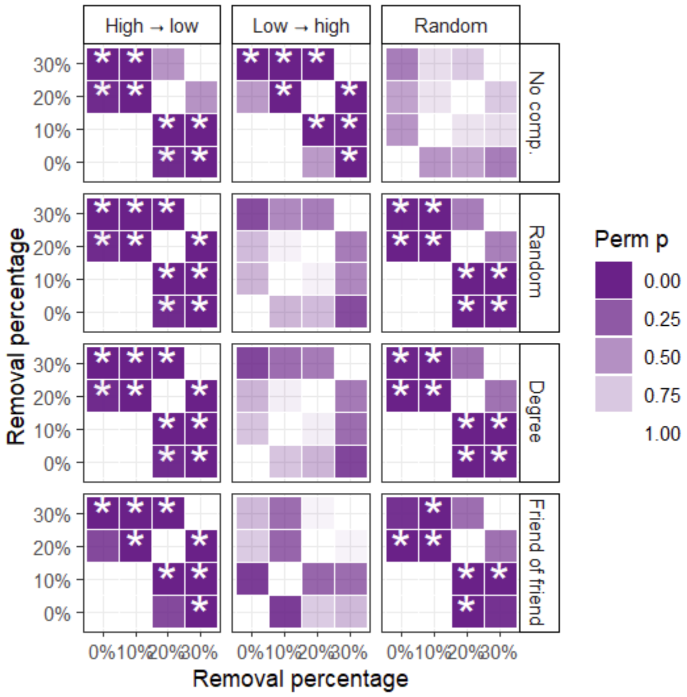

*Figure SI.17: A table of p-values generated by 1000 permutations of compensation type within networks, and calculating resulting percentage changes in clustering coefficient. Stars represent significant (p < 0.05) comparisons.*

*Modularity*

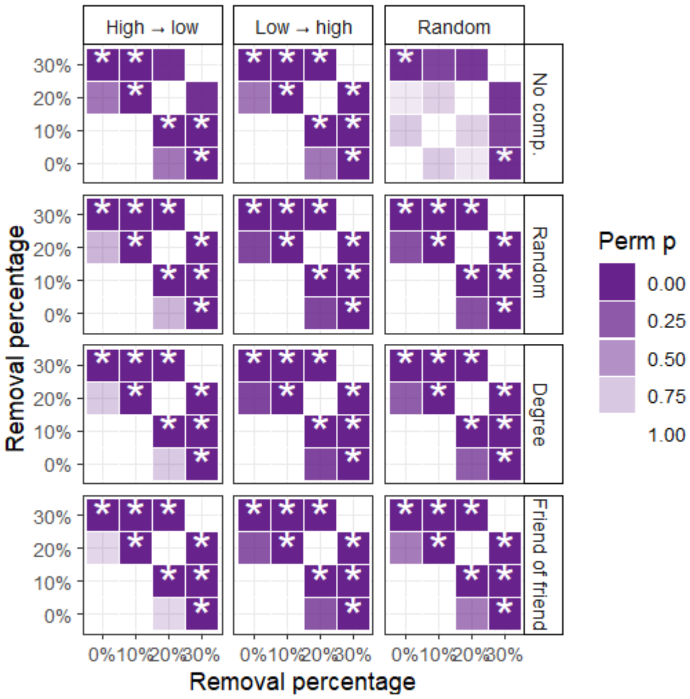

*Figure SI.18: A table of p-values generated by 1000 permutations of compensation type within networks, and calculating resulting percentage changes in modularity. Stars represent significant (p < 0.05) comparisons.*

*Degree heterogeneity*

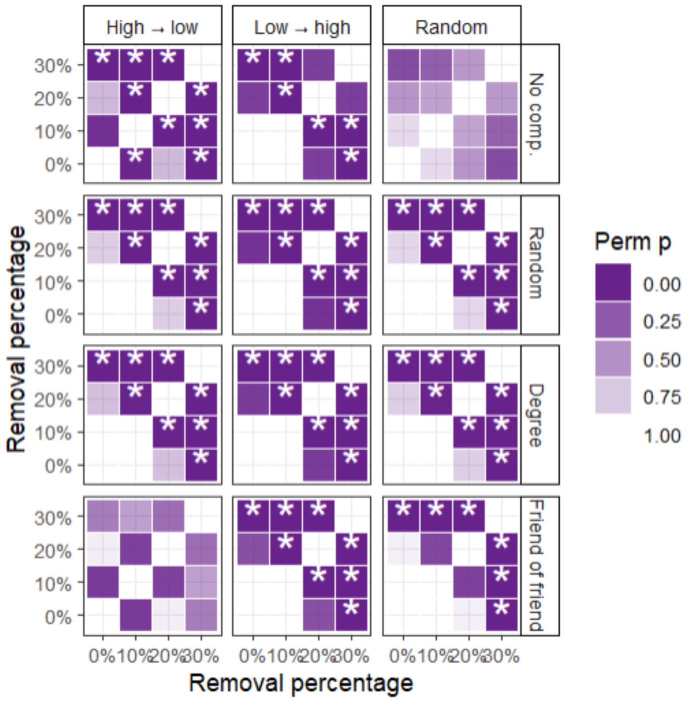

*Figure SI.19: A table of p-values generated by 1000 permutations of compensation type within networks, and calculating resulting percentage changes in degree heterogeneity. Stars represent significant ($p < 0.05$) comparisons.*

*Degree assortativity*

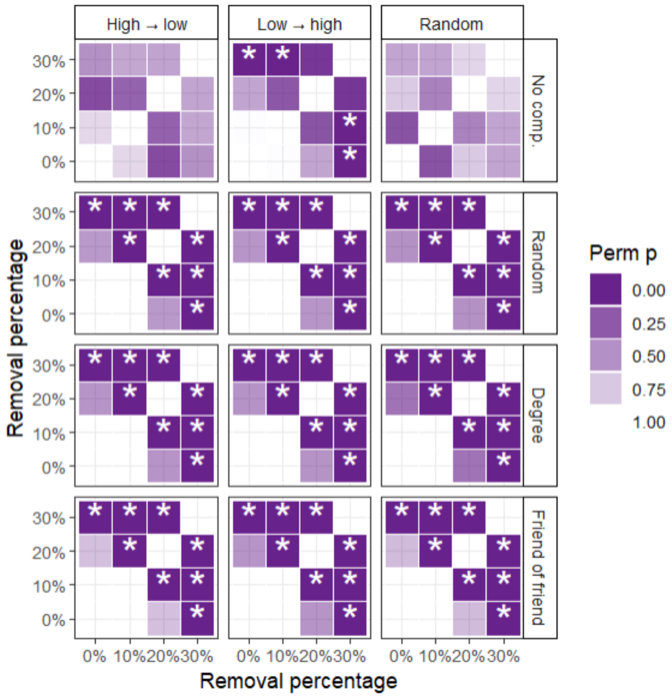

*Figure SI.20: A table of p-values generated by 1000 permutations of compensation type within networks, and calculating resulting percentage changes in degree assortativity. Stars represent significant ($p < 0.05$) comparisons.*

#### Standard deviation

Figure SI.21 shows the range of standard deviations of network results for each metric, in random seed 1. There is no standard deviation for no compensation, as it has no random element (apart from through random removal).

Standard deviation appears to increase for all metrics as removal increases, which matches what we would expect theoretically. Table SI.2 shows how the standard deviation for each metric compares to the range of network mean values for that metric.

*Table SI.2: The network minimum, maximum, and standard deviation values for each of four network metrics.*

| **Coefficient** | **Minimum mean value** | **Maximum mean value** | **Mean SD (% of range)** | **Max SD (%) of range** |
|---|---|---|---|---|
| Clustering | 0 | 1 | 0.045 (4.5%) | 0.465 (46.5%) |
| Modularity | 0 | 0.938 | 0.022 (2.3%) | 0.141 (15.0%) |
| Degree heterogeneity | 0 | 4.58 | 0.035 (0.7%) | 0.157 (3.4%) |
| Degree assortativity | 0 | 1 | 0.042 (4.2%) | 0.234 (23.4%) |

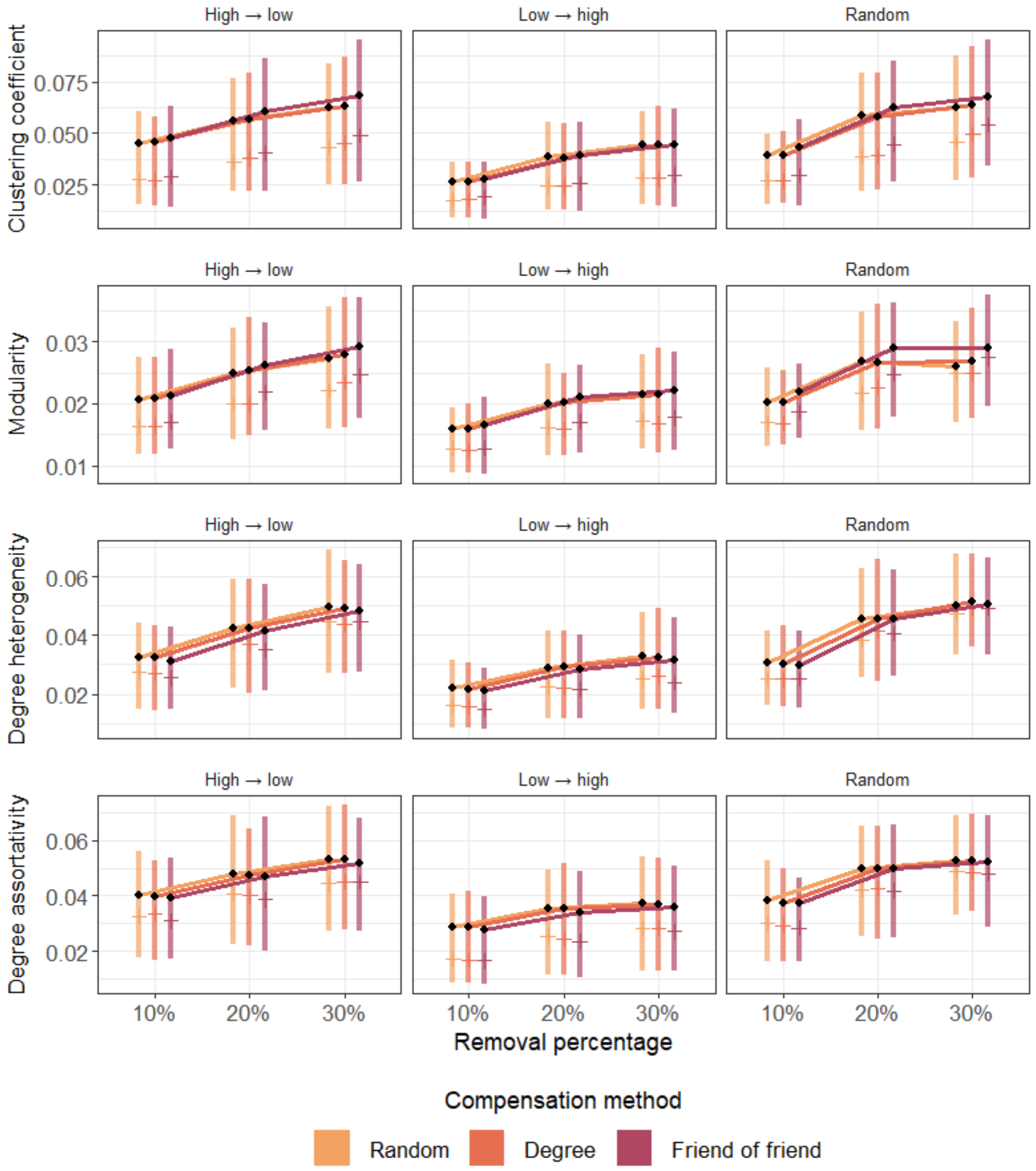

*Figure SI.21: Comparison of standard deviation of network metrics for three removal methods; high-low centrality, low-high centrality and random removal. Compensation method is shown as random compensation (yellow), preserving degree (orange), and bias to friends of friends (red). Lines between black points represent mean values. Line ranges contain 50% of the data, between the 25% and 75% quantile. Small side bars represent median values.*

### Phylogenetic analysis

There were three levels of pseudoreplication within the data - network id, species, and phylogeny. Each network had measures for each of the removal and compensation methods, and some species were represented by multiple networks. Further, certain species were more closely related than others. Figure SI.22 shows a phylogenetic tree of included species, along with the number of networks included from that species. As shown, the dataset is dominated by networks from the field vole *Microtus agrestis*.

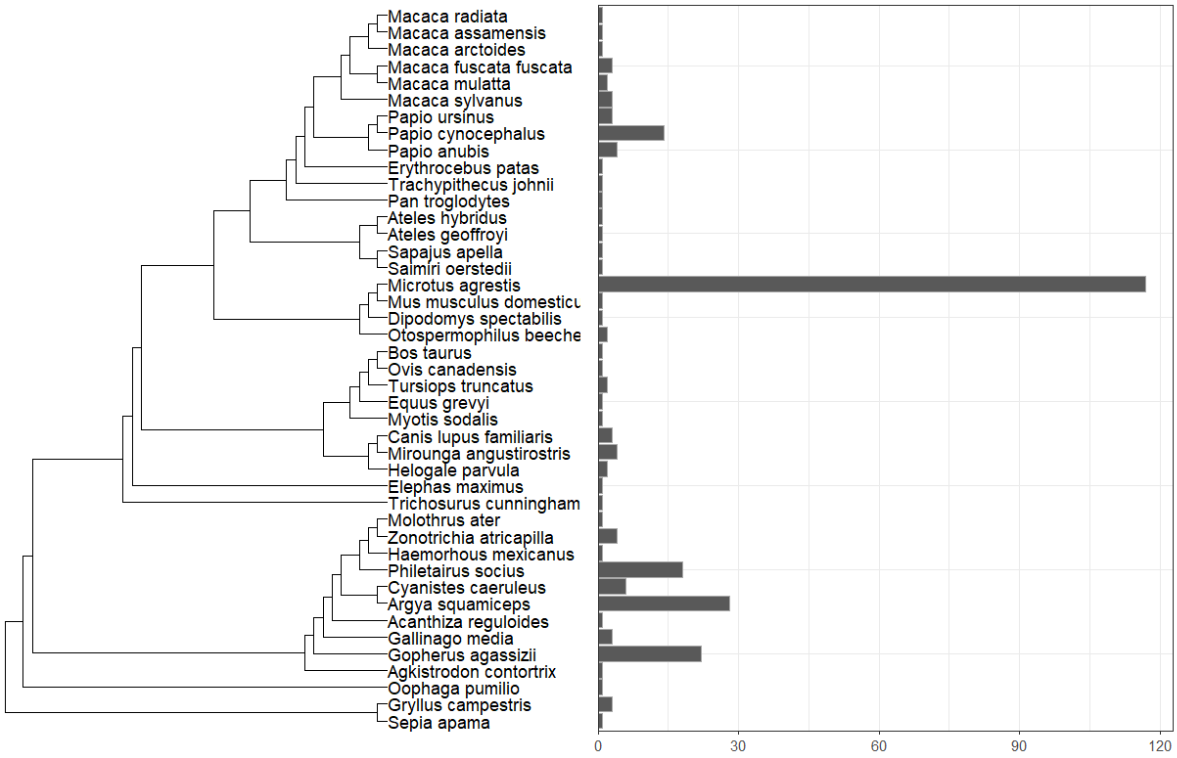

*Figure SI.22: The number of networks associated with each species at 30% removal.*

In order to analyse how phylogenetic pseudoreplication within the dataset explains variation within the data, we created three models of effects with one random effect, each at a different level of phylogenetic variance. This process is explained in more detail in the methods, with the variance explained by the random effect in each of these models presented below (Figure SI.23).

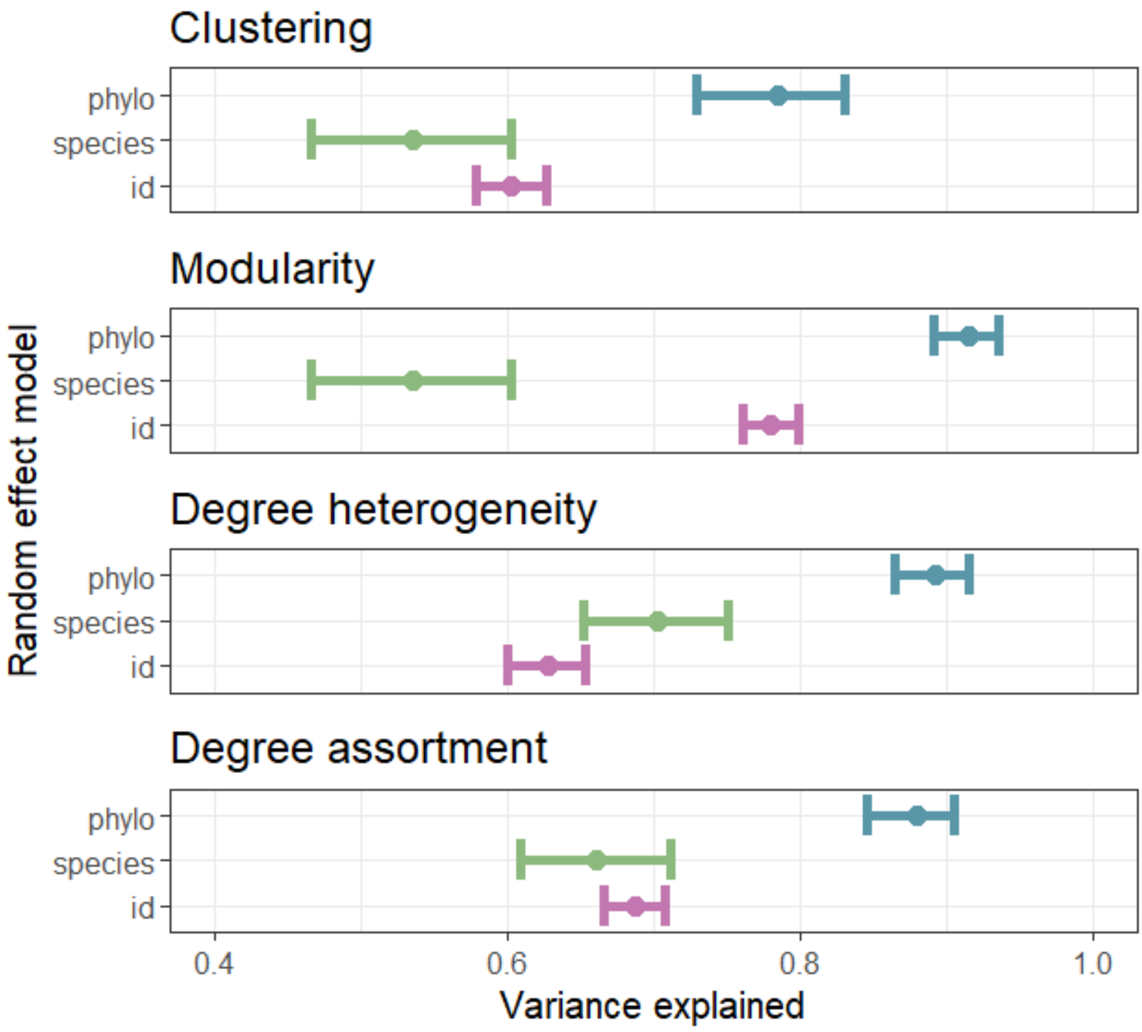

*Figure SI.23: The variance explained by each of three models for four different network metrics. Each model had a different random effect, which is listed on the y-axis. ‘Phylo’ was a phylogenetic covariance matrix of the relatedness of the network species. ‘Species’ was the network species, and ‘id’ was the individual network. Variance explained refers to the proportion of the residual variance explained by this random effect, after controlling for removal method, compensation method, and metric start value.*

Full model results for each random effect structure are presented below (Figure SI.24).

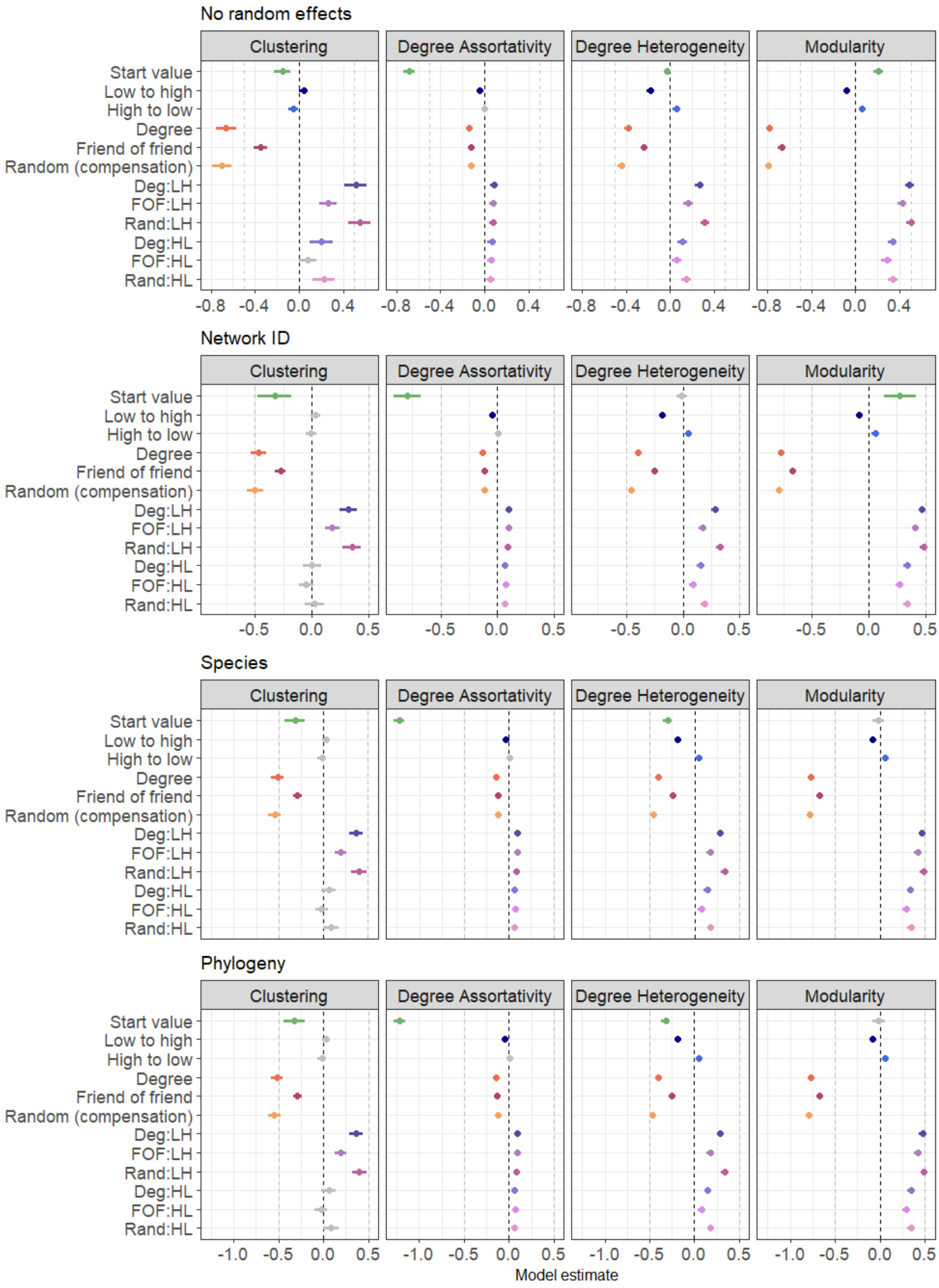

*Figure SI.24: Estimated effect of removal method, compensation method, and start value on the multiplication factor of each metric, estimated from models with random effects accounting for one of four levels of pseudoreplication. Estimates are calculated using a Bayesian Student's t model in brms (Bürkner 2021). Points are point estimates, with lines representing 95% credible intervals. Random removal and no compensation are used as reference levels. Acronyms are as follows: Deg (degree compensation); FOF (Friend of*

*friend compensation); Rand (Random compensation); LH (Low-to-high removal); HL (High-to-low removal). Points coloured in grey have credible intervals that include zero.*

The response variable of these models was multiplication factor (MF), the calculation of which is described in the main text methods. One limitation of this variable is the inability to calculate MF for networks which begin or end with a metric value of 0. Table SI.3 presents the number of networks which were unable to be included in the models for this reason.

*Table SI.3: The number and percentage of networks with an unusable MF value for each metric. MF=NA means a network began at a positive value and decreased to 0 with compensation, or rewiring was not possible in the network, while MF=Inf means a network began at a value of 0 and increased with compensation.*

| **Coefficient** | **Sample size** | **MF NA** | **MF Inf** |
|---|---|---|---|
| Clustering | 3050 | 150 (4.92%) | 174 (5.71%) |
| Modularity | 3050 | 111 (3.64%) | 242 (7.93%) |
| Degree heterogeneity | 3050 | 101 (3.31%) | 12 (0.39%) |
| Degree assortativity | 3050 | 136 (4.46%) | 18 (5.90%) |

Because infinite MF values were generated by start values of 0, these were phylogenetically determined, in that networks from the same species are more likely to have similar network structures. Alternatively, structural similarities between networks of related species could also be due to similarities in data collection methods. As can be seen in Figure SI.25 below, networks with infinite values of MF in clustering tended to also have infinite values in modularity, but this relationship did not hold to degree heterogeneity and degree assortativity. Removed MFs do appear to be more common in particular areas of the phylogenetic tree, suggesting that our analysis may be biased due to our removal of infinite and NA MFs. However, degree heterogeneity and degree assortativity appear to be less biased. The role of phylogeny in our results merits further research, which may disentangle the possible effect of this bias.

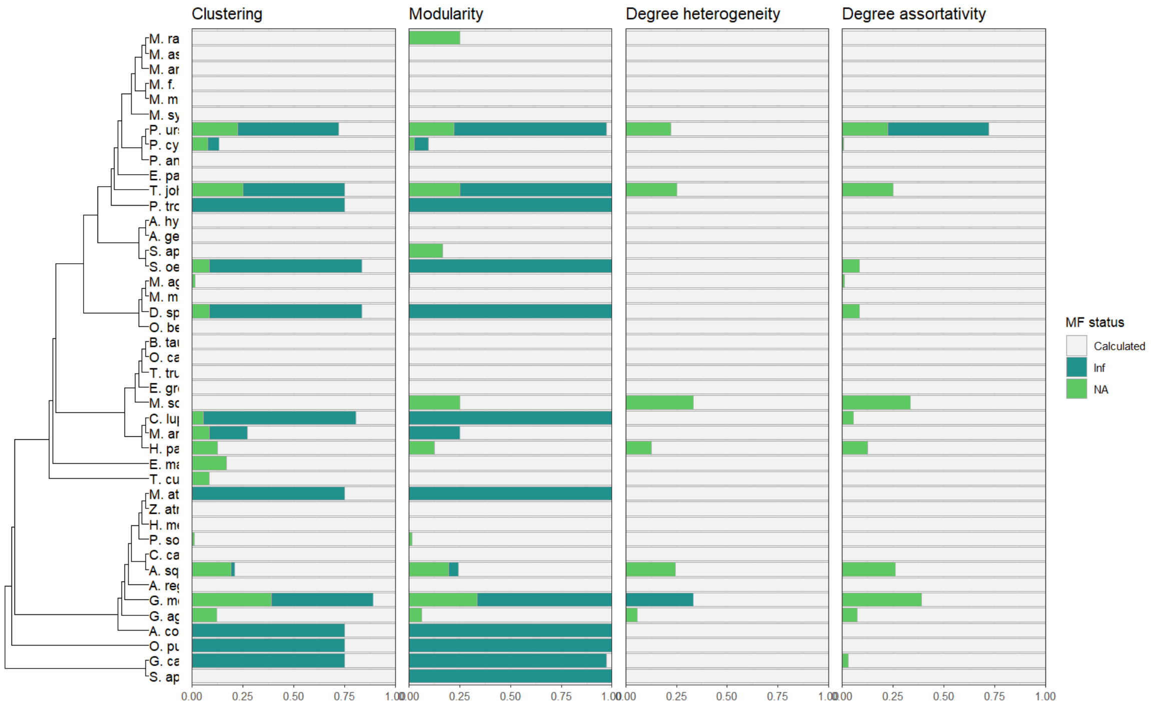

*Figure SI.25: The number of networks where the multiplication factor was able to be calculated, not able to be calculated because metric start value was 0 (Inf) or not able to be calculated because metric value after compensation was 0 (NA). Results are presented by species, across four network metrics.*
